# Predicting phenotype transition probabilities via conditional algorithmic probability approximations

**DOI:** 10.1101/2022.09.21.508902

**Authors:** Kamaludin Dingle, Javor K. Novev, Sebastian E. Ahnert, Ard A. Louis

## Abstract

Unravelling the structure of genotype-phenotype (GP) maps is an important problem in biology. Recently, arguments inspired by algorithmic information theory (AIT) and Kolmogorov complexity have been invoked to uncover *simplicity bias* in GP maps, an exponentially decaying upper bound in phenotype probability with increasing phenotype descriptional complexity. This means that phenotypes with very many genotypes assigned via the GP map must be simple, while complex phenotypes must have few genotypes assigned. Here we use similar arguments to bound the probability *P* (*x* → *y*) that phenotype *x*, upon random genetic mutation, transitions to phenotype *y*. The bound is 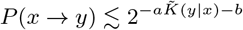, where 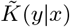 is the estimated conditional complexity of *y* given *x*, quantifying how much extra information is required to make *y* given access to *x*. This upper bound is related to the conditional form of algorithmic probability from AIT. We demonstrate the practical applicability of our derived bound by predicting phenotype transition probabilities (and other related quantities) in simulations of RNA and protein secondary structures. Our work contributes to a general mathematical understanding of GP maps, and may facilitate the prediction of transition probabilities directly from examining phenotype themselves, without utilising detailed knowledge of the GP map.

## I. INTRODUCTION

An important challenge within theoretical biology is understanding the structure of *genotype-phenotype (GP) maps*, which dictate how gene sequences are translated into different biological forms, functions and traits, known as phenotypes. Elucidating GP map structure is essential to a proper understanding of evolution [1] because while random mutations occur at the genetic level, the effects of mutations occur at the level of the phenotype and therefore depend on GP map structure.

Several common properties of GP maps have been identified [2, 3], such as a strong bias in terms of how many genotypes are assigned to each phenotype [4–7] and high degrees of robustness to genetic mutations [8–11]. Significantly, GP map structure has been shown to strongly influence the trajectories and outcomes of evolution: Computer simulations of the evolution of RNA secondary structures (SS) [12, 13], protein complexes [6, 14], genetic circuits [15], among others [16], have shown that even in the presence of natural selection the bias arising from GP map structure can influence and even dominate outcomes. More significantly, for naturally occurring RNA shapes [9, 12, 17–22] and protein quaternary structures [23], the frequency in nature of different molecular shapes can be predicted from GP map biases. The way genotypes are associated to phenotypes via the GP map is also known to limit and constrain evolutionary accessibility of phenotypes [24–26].

Despite the importance of GP maps and the observed common properties across a variety of example maps, the theoretical underpinnings for these observations are not well developed. A recent approach to mathematically studying GP map structure is to use arguments inspired by *algorithmic information theory* [27–29] (AIT), a field of computer science that studies the information content and complexity of discrete patterns, structures, and shapes. Based on these arguments, in refs. [23, 30, 31] it was shown that the estimated information content, or *Kolmogorov complexity*, of a phenotype shape is closely connected to the probability that the shape appears on random sampling of genotypes. Moreover, high-probability phenotype shapes were shown to be simple, and more complex shapes were exponentially less probable, leading to the discovery of *simplicity bias* in GP maps. Interestingly, this complexity approach has enabled predicting the frequency with which biomolecule shapes appear in databases of natural biomolecules [23, 32]. More broadly, many studies have shown that employing AIT as a theoretical framework combined with estimates of Kolmogorov complexity can be fruitful in the natural sciences. Example applications include in thermodynamics [33–35], understanding the regularity of the laws of physics [36], entropy estimation [37, 38], classifying biological structures [39], evolution theory [40, 41], networks [42, 43], in addition to data analysis [44–46] and time series analysis [47, 48], among others [49].

Here, we extend the earlier work on simplicity bias in GP maps by utilising information complexity arguments to predict phenotype transition probabilities: We use AIT as a theoretical framework to derive probability-complexity relations in order to make predictions about the probability *P* (*x* → *y*) that phenotype *y* appears upon the introduction of a single random genetic point mutation in a genotype coding for the phenotype *x*. We show that *P* (*x* → *y*) is fundamentally related to the conditional complexity of *y* given *x*, which measures how much information is required to produce phenotype *y* given access to *x*. We derive an upper bound equation and test it computationally on models of RNA and protein secondary structure, finding good quantitative accuracy.

## II. NULL MODELS FOR TRANSITION PROBABILITIES

### A. Problem set-up

We will focus on GP maps which have some large (finite) number *N*_*g*_ of discrete genotype sequences, a large number of *N*_*p*_ of possible phenotypes, each of which is designable (i.e. has at least one genotype). Further, we will assume that there are many more genotypes than phenotypes, such that 1 ≪ *N*_*p*_ ≪ *N*_*g*_, which is a common property of well-studied GP maps [3]. The GP map will be denoted *f*. The phenotypes are assumed to be some kind of discrete pattern, sequence, or shape, or at least one that could be discretised. For example, a protein quaternary structure can be represented as a discrete graph of nodes [32], and a continuous chemical concentration-time curve can also be discretised in a number of ways [30]. We assume the map is deterministic, such that each genotype maps consistently to only one phenotype.

As an example of such a GP map, we can take the RNA sequence-to-secondary structure map for which a genotype of length *L* featuring four nucleotides (A, U, C, and G) has *N*_*g*_ = 4^*L*^ possible sequences. Also, *N*_*p*_ *∼*1.8^*L*^ [20] so that 1 ≪ *N*_*p*_ ≪ *N*_*g*_ even for modest *L*, and the phenotypes can be represented as discrete sequences because RNA SS are commonly given in dot-bracket form, consisting of a string of *L* symbols defining the bonding pattern of the molecule. In contrast to this RNA example, we are not considering GP maps with only few phenotypes such as whether a patient does or does not have cancer, for which there are only two possible phenotypes, and ‘has cancer’ is not a discrete pattern or shape.

We will write *P* (*x*) for the probability that phenotype *x* appears when uniformly randomly sampling a genotype out of the full collection of *N*_*g*_ genotypes. *P* (*x*) will be called the *global frequency* of *x*. Although the average probability will be 1*/N*_*p*_, possibly for some phenotypes *P* (*x*) ≫ 1*/N*_*p*_ due to bias, and also possibly *P* (*x*) *≪* 1*/N*_*p*_ for some phenotypes. The *neutral set* (NS) of *x* is the set of genotypes that map onto *x*. If we pick one random genotype *g* from the NS of *x*, and make a random single point mutation such that we have a new genotype *g*′, we will call the resulting phenotype *y*. It is possible that phenotypes *x* and *y* are the same or different, because *g*′ may possibly be in the NS of *x*.

If *x* = *y*, then we designate the mutation as neutral. We will define *P* (*x* → *y*) as the transition probability that a randomly selected genotype from the NS of *x*, upon a single random mutation, yields the phenotype *y*. Note that we will still use the word ‘transition’ even if *x* = *y*. A phenotype is called *robust* to mutations if the probability that *y* = *x* is high, i.e. the phenotype typically remains unchanged after a random mutation. The high robustness of phenotypes to genetic mutations is essential to life, and evolution (at least as we know it) would not be able to proceed without it [8, 50]. The origin of high robustness has been seen as something of a mystery, however [50]. Additionally, it has been noticed in several GP maps that robustness scales with the logarithm of the global frequency of a phenotype, as opposed to scaling with the global frequency itself which would be expected from a random null model. The cause of this general logarithmic scaling is presently not fully explained, although an abstract model of GP maps has been used to study this property [51]. In a future study we intend to look in detail at robustness.

The problem of estimating *P* (*x* → *y*) is the main focus of the work, and in particular relating this quantity to the relative information contents of *x* and *y*.

### B. Null models of transition probability

How can we estimate the transition probability *P* (*x* → *y*)? If we have access to data recording the frequency of transitions in simulations, then we could directly estimate *P* (*x* → *y*) from those data by counting the number of times *x* transitioned to *y* as a fraction of all transitions starting with *x*. It may also be possible to arrive at an estimate of *P* (*x* → *y*) by examining details of the map and the particular phenotype *x*. However, what we are interested in here is a general method for predicting *P* (*x* → *y*), that does not rely on using past frequency data, or details of the map. Indeed, we are interested in general properties of GP maps which will both help to develop a theory of GP maps, and also be useful for other maps for which we neither have data nor a clear understanding of exactly how the phenotype arises from the genotypes. In a sense we are interested in *a priori* prediction for *P* (*x* → *y*) which is based only on the patterns in *x* and *y*. At first sight, this goal may not appear possible, because *P* (*x* → *y*) *will* depend on the details of the map. However, we will argue in this work that even without knowing details of the map, and without recourse to historical frequency data, non-trivial predictions for the transition probabilities can be derived. In another sense, we are interested in establishing a good null model for *P* (*x* → *y*), which could serve as a starting point for predictions about transitions. In this connection, we now consider some possible null models and weigh up their merits.

Perhaps the simplest null model expectation is that the transition probability is

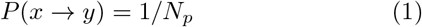

which corresponds to a maximum entropy estimate, assigning a uniform probability to each possible outcome *y*. However, this has a limitation which is that a common property of GP maps is bias (described above), and so it seems reasonable to expect some degree of non-uniformity in *P* (*x* → *y*). Further, the high levels of robustness discussed above do not accord with this uniform distribution model. From here onwards, we will assume that the distribution *P* (*x* → *y*) over the possible values of *y* is strongly non-uniform (biased).

Another simple null model for *P* (*x* → *y*) was proposed in ref. [13]

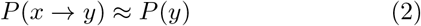

for *y* ≠ *x*. This null model prediction is correct if genotypes are randomly assigned to phenotypes, with no correlations between genotypes or phenotypes. While the approximation in Eq. (2) is straightforward and was observed to be quite accurate on average [13], it also has limitations. Firstly, as pointed out in ref. [30] many GP maps have fixed and somewhat simple rule sets by which genotypes are assigned to phenotypes (technically, they are *O*(1) complexity maps). Hence, these maps do not randomly assign genotypes, but assign them with a definite structure and pattern, which is likely to produce some clear patterns in genotype architecture. Secondly, it is intuitively reasonable that phenotype *x* will transition to some *y* which is similar or even the same as *x*. The logic being that one single point mutation represents a small change to the genotype, and consequently a small change to the phenotype appears to be a rational null assumption. Of course this assumption that GP maps are roughly ‘continuous’ in the mathematical sense of the word does not always hold because some (well-chosen) mutations may drastically change the phenotype, but nonetheless the assumption has intuitive appeal and may typically hold. Greenbury et al [50] have also suggested that transitions are more likely to be to similar phenotypes (with the caveat that it must be possible for enough genotypes to be sampled), arguing via genetic correlations in GP maps. Hence Eq. (2) has limitations as a null model.

In order to improve on Eq. (2), we would like to incorporate the ‘similar phenotypes’ notion in a formal way, which will lead to a new null model that we propose. To do this we first need to survey some pertinent theoretical background.

## II. ALGORITHMIC INFORMATION THEORY

### A. Kolmogorov complexity

Developed within theoretical computer science, *algorithmic information theory* [27–29] (AIT) connects computation, computability theory, and information theory. The central quantity of AIT is *Kolmogorov complexity, K*(*x*), which measures the complexity of an individual object *x* as the amount of information required to describe or generate *x*. More formally, the Kolmogorov complexity *K*_*U*_ (*x*) of a string *x* with respect to a universal Turing machine [52] (UTM) *U*, is defined [27–29] as

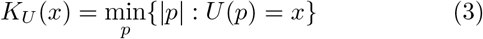

where *p* is a binary program for a prefix (optimal) UTM *U*, and *p* indicates the length of the (halting) program |*p*| in bits. Due to the invariance theorem [49] for any two optimal UTMs *U* and *V, K*_*U*_ (*x*) = *K*_*V*_ (*x*) + *O*(1) so that the complexity of *x* is independent of the machine, up to additive constants. Hence we conventionally drop the subscript *U* in *K*_*U*_ (*x*), and speak of ‘the’ Kolmogorov complexity *K*(*x*). Despite being a fairly natural quantity, *K*(*x*) is uncomputable, meaning that there cannot exist a general algorithm that for any arbitrary string returns the value of *K*(*x*). Informally, *K*(*x*) can be defined as the length of a shortest program that produces *x*, or simply as the size in bits of the compressed version of *x*. If *x* contains repeating patterns like *x* = 1010101010101010 then it is easy to compress, and hence *K*(*x*) will be small. On the other hand, a randomly generated bit string of length *n* is highly unlikely to contain any significant patterns, and hence can only be described via specifying each bit separately without any compression, so that *K*(*x*) ≈ *n* bits. *K*(*x*) is also known as *descriptional complexity, algorithmic complexity*, and *program-size complexity*, each of which highlight the idea that *K*(*x*) measures the amount of information required to describe or generate *x* precisely and unambiguously.

An important quantity for our present investigation is the *conditional complexity, K*(*y*|*x*), defined as

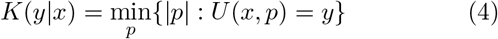

i.e., the minimum length of a program *p* such that a UTM *U* generates string *y*, given *x* and *p* as an input. Less formally, *K*(*y* | *x*) quantifies how many bits of information are required to generate *y*, given that we have access to *x*.

More details and technicalities can be found in standard AIT references [49, 53–55] and a book aimed at natural scientists [56].

### B. Algorithmic probability

In AIT, Levin’s [57] coding theorem establishes a fundamental connection between *K*(*x*) and probability predictions. Building on Solomonoff’s discovery of *algorithmic probability* [27, 58], Levin’s coding theorem [57] states that

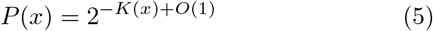

where *P* (*x*) is the probability that (prefix optimal) UTM *U* generates output string *x* on being fed random bits as a program. Thus, high-complexity outputs have exponentially low probability, and simple outputs must have high probability. *P* (*x*) is also known as the *algorithmic probability* of *x*.

The *conditional coding theorem* [59] states that the probability *P* (*y* | *x*) of generating string *y* with UTM *U* given access to string *x* as side information is

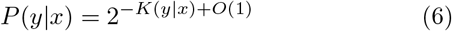

Notice that outputs with high probability here must have low conditional complexity *K*(*y* |*x*). To have *K*(*y* | *x*) low means that either *y* is simple itself, or it is similar to *x*. To see this, consider that if *y* is simple, then *K*(*y*) is low, then so too is *K*(*y*|*x*), hence *P* (*y*|*x*) is high. Also, if *y* is similar to *x* — for example they share common subsequences — then *K*(*y*|*x*) will be low, and *P* (*y*|*x*) will be high.

### C. Simplicity bias

Eq. (5) as well as many other AIT results cannot be straightforwardly applied to typical natural systems of interest in engineering and sciences, due to the fact that: (1) Kolmogorov complexity is uncomputable and so cannot be calculated exactly; (2) the theory is asymptotic, valid only up to *O*(1) terms; (3) the theory is largely based on UTMs, which are seldom present in nature; and (4) the coding theorem assumes infinite purely uniform random programs, which do not exist in nature.

Despite these points, several lines of reasoning motivate using AIT to make predictions while being aware of the limitations of this practice. We call this kind of theoretical work ‘AIT-inspired’ arguments. See Supplementary Information (SI) III (A & B) for more discussion on this.

Adopting the methodology of AIT-inspired arguments, Dingle et al [30] studied coding theorem-like behaviour and algorithmic probability for (computable) real-world inputoutput maps. This led to their observation of *simplicity bias* (SB), governed by the equation

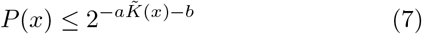

where *P* (*x*) is the (computable) probability of observing output *x* on random choice of inputs, and 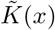 is the approximate Kolmogorov complexity of the output *x*. In words, SB means complex outputs from input-output maps have lower probabilities, and high probability outputs are simpler. The constants *a >* 0 and *b* can be fit with little sampling and often even predicted without recourse to sampling [30].

Examples of systems exhibiting SB are wide-ranging, and include molecular shapes such as protein structures and RNA [23], outputs from finite state machines [31], as well as models of financial market time series and ODE systems [30], among others. A full understanding of exactly which systems will, and will not, show SB is still lacking, but the phenomenon is expected to appear in a wide class of input-output maps, under fairly general conditions. See SI III (C) for more on this.

## IV. SIMPLICITY BIAS IN TRANSITIONS

### A. Simplicity bias: conditional form

Just as for the original coding theorem, the conditional coding theorem in Eq. (6) cannot be directly applied to practical real-world systems, such as making estimates for phenotype transition probabilities. So we derive (SI III (D & E)) a conditional form of the simplicity bias equation Eq. (7), which we subsequently apply to phenotype transition probabilities: The conditional form is

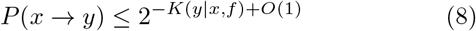

### B. Complexity of the GP map

The complexity of the GP map *f* is an important quantity. If the map *f* is allowed to have a high complexity value, then *f* could be chosen such that *P* (*x* → *y*) takes arbitrary values, and hence it will be very hard to predict transition probabilities. Fortunately, many GP maps are not random, but in fact have simple (low-complexity) fixed rule-sets for determining how genotypes are assigned to phenotypes [30]. See SI III (F) for more details.

If we restrict our attention to GP maps for which *f* is of fixed complexity, i.e. *K*(*f*) = *O*(1), then this means that *K*(*y*|*x, f*) ≈ *K*(*y*|*x*) so that Eq. (8) becomes

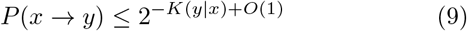

and we see that this upper bound depends only on the phenotypes *x* and *y*. So the complexity of the map *f* is an important quantity which either does or does not allow predictions to be made just using conditional complexities.

### C. Approximation of the upper bound

Because Kolmogorov complexity is uncomputable, in practice we use approximations, such as real-world compression algorithms [49] (See also SI III (B) for more on this). Following the approximation and scaling arguments of ref. [30], we can write an approximate form of Eq. (9)

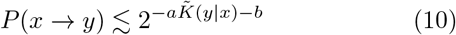

which is a weaker form of the full AIT conditional coding theorem [59] given in Eq. (6). The term 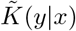 is an approximation of the conditional Kolmogorov complexity *K*(*y* | *x*), which we will calculate according to the Lempel-Ziv [60] complexity estimate used earlier [30, 31] and also scale the complexity values so that 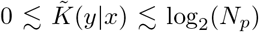 as described in the methods in SI I (A). To estimate the conditional complexity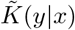, we employ the approximation (as used earlier [61]) that

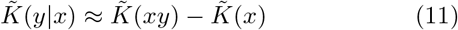

where 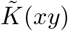 is the compressed length of the concatenation of strings *x* and *y*. For example, if *x* =`ABC` and *y* =`XY` then *xy* =`ABCXY`. Note that for true prefix Kolmogorov complexity the relation *K*(*y* | *x*) ≈ *K*(*x, y*) −*K*(*x*) only holds to within logarithmic terms [49], but that is close enough for our purposes. Note that the terms *K*(*x, y*) and *K*(*xy*) are quantitatively very close, especially if the lengths of *x* and *y* are the same. Hence we make the approximation that they are equal.

The constant *a* may depend on the map, but not on the phenotype *x*. If the complexity 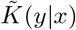 is scaled properly (SI I(A)), then *a* = 1 is the default prediction. Otherwise, *a* might have to be fit to the data. The main requirement for scaling properly is having a reasonably accurate estimate of the number *N*_*y*_(*x*) of phenotypes *y* such that *P* (*x* → *y*) *>* 0, i.e., the number of accessible phenotypes via a single point mutation from *x*. If *N*_*y*_(*x*) is known a priori or can be estimated a priori, then *N*_*y*_(*x*) can be used for a priori prediction of *P* (*x* → *y*). Otherwise, if random genotype sampling is employed, then simply counting the number of different *y* phenotypes observed in sampling is one way to estimate *N*_*y*_(*x*). Naturally, this counting method will be more accurate for larger samples, and may produce very low underestimates of *N*_*y*_(*x*) for small sample sizes.

The constant *b* has default value *b* = 0 [30], but can also be found by fitting to the data if necessary. Looking at the examples of simplicity bias presented in the literature to date, it appears that *b* = 0 often works very well.

It follows that in practice, provided some reasonable estimate of the number of accessible phenotypes *N*_*y*_(*x*) is known and hence complexities are scaled well, then Eq. (10) reduces further to the practically applicable relation

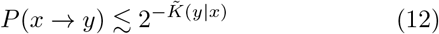

allowing transition probabilities to be made just based on phenotype conditional complexities.

### D. The bound is close with high probability

Based on the arguments in [31], we expect the upper bound Eq. (10) (and also Eq. (12)) to be tight for *x, y* pairs which are generated by random genotypes. That is, for a phenotype *x* generated by a random genotype, and *y* subsequently arising from a random mutation, we expect 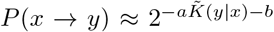 with high probability, as opposed to the right-hand side being only a loose upper bound.

On the other hand, the ubiquity of low-complexity, low-probability outputs [31, 62] suggests that for many *y* we may have 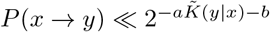. Such phenotypes *y* are those that have low conditional complexity, yet at the same time appear with low probability due to map-specific constraints and biases. See ref. [62] for an in-depth discussion of this low-complexity, low-probability phenomenon.

### E. Size of the genotype alphabet and number of mutations

For the bound of Eq. (10) to have stronger predictive value on point mutations, we suggest that the size of the genotype alphabet *α* should be small. This is not a very onerous condition, and is in fact quite naturally satisfied. Further, the number of mutations should be *≈*1. See SI III (G) for more on these conditions.

### F. When is *P* (*y*) a good predictor of *P* (*x* → *y*)?

From AIT we know that almost all pairs of phenotypes *x* and *y* share almost no information, in other words *K*(*x, y*) ≈ *K*(*x*) + *K*(*y*), so that

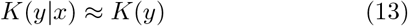

From this we can infer that for almost all pairs of phenotypes *x* and *y* the conditional complexity *K*(*y* | *x*) in Eq. (8) can be replaced with just *K*(*y*), and so the equation becomes

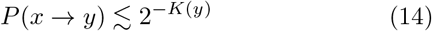

for almost all *y*.

The preceding argument suggests that for most outputs *y*, the phenotype *x* may be largely irrelevant in estimating the probabilities *P* (*x* → *y*). However, this statement comes with the caveat that nearly all the probability mass is likely to be associated to only a small fraction of the possible outputs, those for which *K*(*y* | *x*) is low. See SI III (H) for more discussion and details.

### G. Predicting which of *P* (*x* → *y*_*i*_) or *P* (*x* → *y*_*j*_) is higher

Another quantity that may be predicted using the preceding theory is the ratio of probabilities for transitioning from one phenotype to different alternative phenotypes. In this section, we describe a method for such predictions, which we test numerically below.

Call *y*_*i*_ the resulting phenotype after a single point mutation to a randomly chosen genotype in the NS of *x*. Call *y*_*j*_ the resulting phenotype after a single point mutation to another independently chosen random genotype, also in the NS of *x*. We can use the preceding theory to predict which of the two phenotypes *y*_*i*_ and *y*_*j*_ has a higher probability directly from complexity estimates. This is interesting because it is often valuable to know whether *P* (*x* → *y*_*i*_) *> P* (*x* → *y*_*j*_) or *P* (*x* → *y*_*i*_) *< P* (*x* → *y*_*j*_), rather than trying to guess the exact values of *P* (*x* → *y*_*i*_) and *P* (*x* → *y*_*j*_). Fortunately, the constants *a* and *b* are not required for predicting this via Eq. (10) because only the relative value of 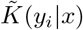 and 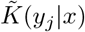 determines whether 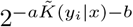 or 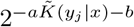 is larger. So even if we could not estimate *a* or *b* accurately, we could still make a prediction about which phenotype is more likely to arise through a point mutation of *x*. See SI I (B) for more details.

### H. Predicting *P* (*x* → *y*_*i*_)*/P* (*x* → *y*_*j*_) ratios

Beyond predicting which has higher probability, we can also try to predict the value of the ratio of probabilities of transitioning from one phenotype to different alternative phenotypes. The ratio is also related to how confident we are in predicting which of the probabilities *P* (*x* → *y*_*i*_) and *P* (*x* → *y*_*j*_) is higher: a higher ratio means more confidence in the prediction.

Because *y*_*i*_ and *y*_*j*_ are both randomly generated, we expect both *P* (*x* → *y*_*i*_) and *P* (*x* → *y*_*j*_) to be close to the bound of Eq. (10), with high probability [30, 31]. Therefore we can use an approximate equality assumption to predict the ratio as follows:

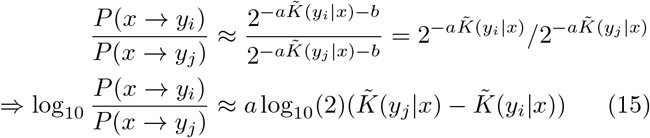

notice that *b* is irrelevant, so that even if *b* is not known, the prediction is unaffected. Recalling from above that we set *a* = 1 yields 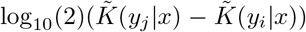 as the predictor for the log_10_ ratio of the probabilities. If the scaling is not done correctly, then *a* ≠ 1 and therefore the predictor will not be as accurate, but instead off by a constant factor. In this case, we still have a relative measure of how confident we are about the prediction, where larger values of 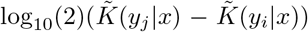 are associated with higher-confidence predictions.

## I. Distribution of conditional complexities

Eq. (10) states that higher-probability phenotypes must have low conditional complexity, so in this sense we have derived a ‘conditional simplicity bias’. A related but different question is, upon choosing a random genotype in the NS of *x*, and introducing a random mutation, what conditional complexity value is most likely? More generally, what kind of distribution should we expect for *K*(*y* | *x*)? This is not a trivial question, because on the one hand, the upper bound on transition probabilities decays exponentially with increasing *K*(*y* | *x*), which would suggest that lower complexities are more likely. On the other hand, from AIT we expect the number of patterns with higher complexity to grow exponentially with increasing complexity, and hence increase with *K*(*y* | *x*), which would suggest that higher complexities are more likely. However, how the actual number of conditional complexities grows for a specific GP map system may not reflect the AIT expectation exactly.

In ref. [23] (see also [48]) it was argued that to a first approximation, these two exponential trends should cancel each other out, leading to a ‘flat’ complexity distribution. Based on these arguments, we suggest that perhaps a flat distribution will also be seen for *K*(*y* | *x*). Predicting the distribution of complexities is somewhat difficult, due to the fact that for a ‘flat’ distribution the two exponential trends must precisely cancel out, while exponential trends can easily magnify small errors in approximation. The distribution of complexities should be investigated in future work.

## V. EMPIRICAL PHENOTYPE TRANSITION PROBABILITIES

### A. Main predictions

Before analysing some example GP maps, we recap the main predictions and equations discussed in this work:

a. (a) High transition probabilities *P* (*x* → *y*) will be associated to either phenotypes which are similar to *x*, or are very simple.
b. Transition probabilities will conform to the upper bound 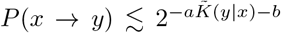 as in Eq. (10), with *a* = 1 and *b* = 0 as default expectations.
c. Some phenotypes will have probabilities close to the upper bound, while many may be low-complexity, low-probability outputs far below the bound.
d. For most phenotypes *y, K*(*y* | *x*) ≈ *K*(*y*) and especially if *K*(*x*) is low, but *K*(*y* | *x*) ≪ *K*(*y*) for some *y*.
e. We can predict which of two phenotypes is more likely to arise just by comparing their conditional complexities, betting on the simpler one having higher probability.
f. We can predict the actual ratio of these two probabilities. In the following sections, we computationally test the applicability of these predictions to RNA and protein SS.

### B. Computational experiment: RNA secondary structure

RNA are important and versatile biomolecules that act as information carriers, catalysts, and structural components, among other roles. Similarly to DNA, RNA is comprised of a sequence of nucleotides, which can contain four possible nucleobases, A, U, C, and G. RNA molecules typically fold up into well-defined SS, which denote bonding patterns in the molecule. The SS shape is determined by the underlying sequence, but at the same time the sequence-to-SS map is highly redundant with many different sequences adopting the same SS. RNA SS has been very well studied in biophysics and computational biology because it is a biologically relevant system, but at the same time fast and accurate computational algorithms exist for predicting SS from sequences. Here we use the popular Vienna RNA [63] package for predicting SS, which utilizes a thermodynamics-based algorithm.

To test our predictions, we randomly generated an RNA sequence of length *L* = 40 nucleotides, and computationally predicted its SS .(((…)))..((((.((..((….)).))))))…., which we will denote by *x*. We chose *L* = 40 because computational efficiency requires a comparatively short sequence, but on the other hand complexity-probability connections are more pronounced for longer RNA [30]. In order to estimate *P* (*x* → *y*), we need to generate a large sample of sequences within the neutral space of *x*, that is, different sequences that each have *x* as their SS. To generate the sequences, we used the site-scanning method of [64]. Afterwards, we introduced a single random point mutation for each neutral sequence, and recorded the resulting SS. More details are given in the methods described in SI I(C).

Figure 1(a) shows the highest-probability SS for each conditional complexity value found in the dataset generated in this way, as well as the estimated probability of transitioning from the starting phenotype *x* to all of these alternative phenotypes. In Figure 1(b) we see that, as expected, there is strong bias in transition probabilities with a decay in the upper bound to *P* (*x* → *y*). The black line is a fit to the data (*a* = 1.1 and *b* = 0) added to highlight the upper bound. Figure 1(b) also shows the predicted upper bound (red line) based on *a* = 1 and *b* = 0; this prediction is impressive, given that it is based on just the output complexities themselves. Note that there are several low-probability structures for which our measure 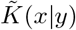 gives zero, even though they are slightly different than the reference structure. This is most likely caused by two effects. Firstly, the lack of very fine resolution in our approximate complexity measure, and second, by our neglecting the O(1) terms. Panel (c) presents the same data as Figure 1(b), except that the horizontal axis is 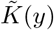 instead of 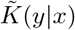, and it is apparent that 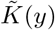 does not provide a good predictor of the probabilities. This demonstrates that the conditional complexities are needed, not just the complexity 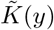. Figure 1(d) is a scatter plot of 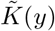 vs 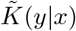 and, as expected, there is a linear (but noisy) correlation between these two quantities (Pearson *r* = 0.64, p-value*<* 10^−6^). It is also interesting to see that, as expected, a small number of phenotypes have conditional complexities 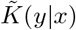 much lower than the unconditional complexity 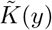, and those tend to be the higher-probability outputs (as can be seen from the colouring of the datapoints by log probability).

**FIG. 1.**
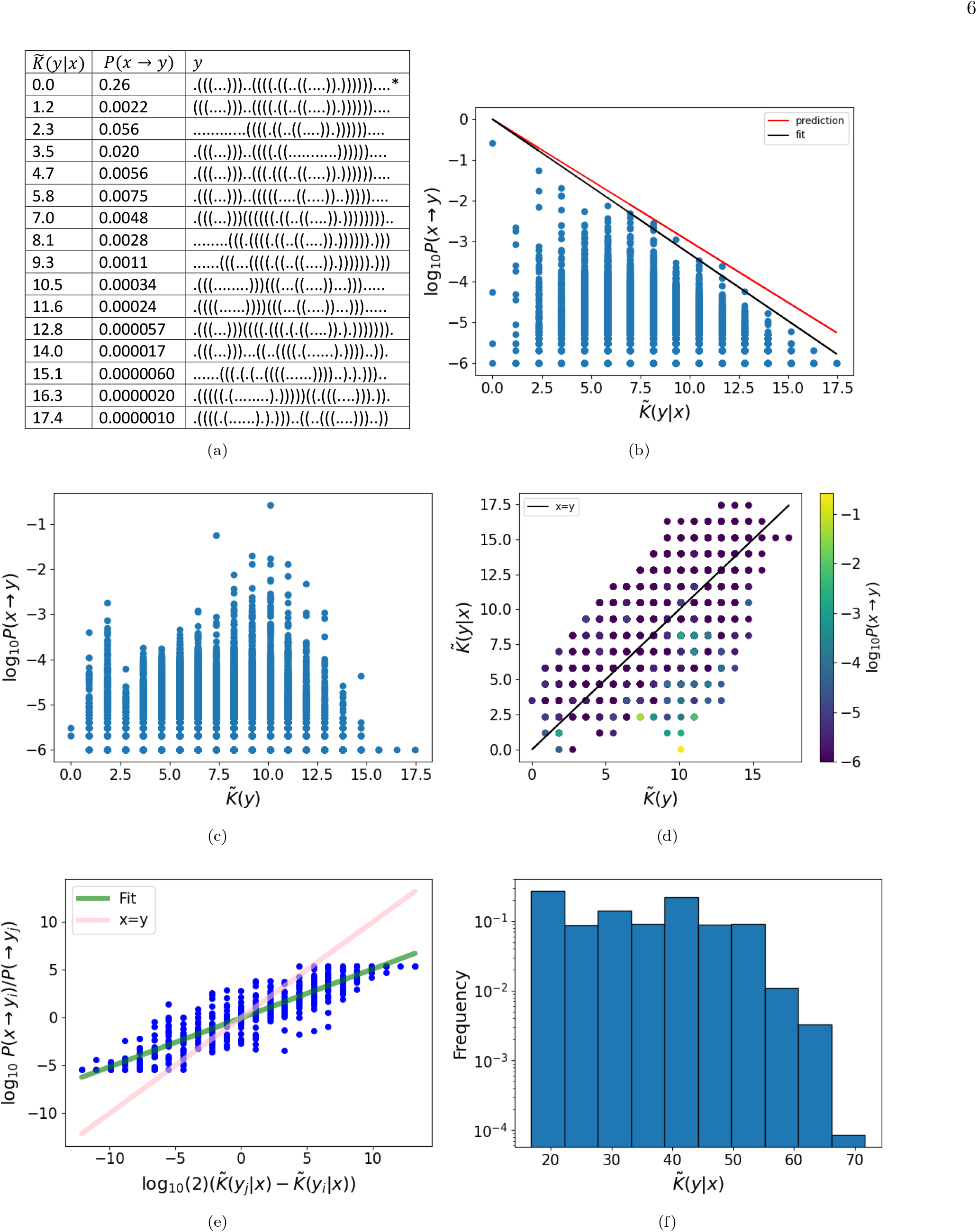
RNA secondary structure transition probabilities for a sequence of length *L* = 40 nucleotides. The starting phenotype is *x* =.(((…)))..((((.((..((….)).)))))). and transitions result from choosing random genotypes in the NS of *x*, and introducing a single random mutation to each genotype. (a) Table illustrating the highest probability SS for each conditional complexity value. The starting phenotype *x* is marked with an asterisk (*), and *P* (*x* → *y*) is just the robustness since *x* = *y*. (b) Transition probabilities *P* (*x* → *y*) decrease exponentially with increasing conditional complexity 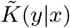, upper bound of Eq. (10) depicted in black. The highest probability SS is the same phenotype as *x*. The predicted upper bound (red) and fitted bound (black) are close. (c) The unconditional complexity 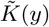 does not predict the transition probabilities well. (d) 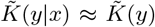 for most *y*, leading to a positive linear correlation between values, cf. Eq. (13). (e) Ratios of probabilities correlate strongly with differences in conditional complexity, partially according with Eq. (15). (f) The histogram of conditional complexity values shows a roughly flat distribution (on a log scale), but with some slight bias towards simplicity.

Turning to the prediction of which of two phenotypes has higher probability, we find that with probability-weighted sampling the accuracy is a striking 86%, and with uniform sampling, the rate is still impressive at 79%. (Recall that probability-weighted sampling refers to when genotypes are uniformly randomly sampled, and hence phenotypes appear with frequencies according to their probabilities. Uniform sampling refers to when each phenotype is sampled with equal probability.) Extending this, Figure 1(e) depicts not just predictions of which is higher, but of the ratios themselves. Although the fit does not match the *x* = *y* prediction line, there is nonetheless a strong correlation (Pearson *r* = 0.91, p-value*<* 10^−6^) between the ratios of the probabilities and the differences in complexities. This means that although the slope is not very well predicted, the expected close connection between the complexity values and the probabilities holds. Recall that the inaccuracy in the slope predictions for Figures 1(b) and (e) results primarily from a lack of precision in estimating the value of *a*, which depends on knowing the number of possible phenotypes *y* such that *P* (*x* → *y*) *>* 0. The fact that the actual slope is flatter than the predicted one is presumably due to the following: The value of 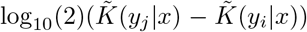 will be large when *y*_*j*_ is (conditionally) complex and *y*_*i*_ is simple. Therefore *y*_*j*_ is unlikely to be far from the upper bound, while very simple phenotypes can be very far from the upper bound, cf. the low-complexity, low-probability phenomenon [31, 62]. Hence the value of *P* (*x* → *y*_*i*_)*/P* (*x* → *y*_*j*_) is likely to be an underestimate, rather than an overestimate, which will tend to make the slope flatter.

Finally, Figure 1(f) presents a histogram of conditional complexities, showing that their distribution is roughly flat on a log scale as tentatively predicted, but it also exhibits some mild bias towards lower-complexity phenotypes.

In SI II we numerically study the impact of the complexity of the starting phenotype *x*, showing that the mean value in the difference between 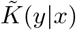 and 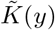 is small for simple *x* as expected, and grows for more complex *x*. Further in SI IV we provide an additional example RNA plot, depicting results similar to those in Figure 1.

### C. Computational experiment: protein secondary structure

Proteins are biomolecules that form the fundamental building blocks of organisms. A protein is comprised of one or more macromolecular chains that in living organisms typically contain 20 types of amino acid residues. Similarly to RNA, a protein will fold into a particular spatial structure, which is determined by the specific amino acid sequence. There is redundancy in that many different sequences can have the same fold, see e.g. [66]. The overall 3D arrangement of a protein’s polypeptide chain in space is known as its tertiary structure, while protein SS refers to the local conformation of the polypeptide backbone. SS is key to protein folding [67] and the genotype-to-phenotype map between primary and secondary structure has not received much attention in the literature to date. At the level of detail we are concerned with, determining a protein’s SS is equivalent to specifying whether each amino acid residue in the chain is involved in a coil (C), sheet (E), or helix (H) structure. Hence, the SS of a protein of length *L* is also a sequence of length *L*, but with only three possible letters (C, E, H) at each site.

Predicting the full 3D structure of a protein was until recently an open problem, but it is now feasible via machine learning algorithms such as AlphaFold [68]. However, it remains very computationally taxing and potentially unreliable for sequences not related to the natural sequences used to train the underlying machine learning algorithm. In contrast, the machine learning-based Porter 5 algorithm provides accurate and relatively fast predictions of protein SS [69]. Predicting the structure of mutants remains challenging both for algorithms such as AlphaFold [70] and for SS ones like Porter 5. Here we make the implicit assumption that Porter 5 captures the large-scale properties of the mapping between protein primary and secondary structure sufficiently accurately that our computational study grants insight into the system. Such insight is particularly valuable considering that it is impractical to survey a similar number of mutants experimentally.

Like with the RNA example, choosing the length *L* of the protein under study is a balance between computational cost and accuracy: If longer proteins are used, then the number of possible SS grows exponentially, making it hard to get accurate estimates of probabilities as the latter require the same SS to appear multiple times. On the other hand, very short proteins are less biologically relevant, and the complexity measures and theory are expected to work worse for them than for longer sequences. Here we balance these considerations by studying a complementarity-determining region (CDR) with *L* = 20, specifically CDRH3 from the heavy chain of a human monoclonal SARS-CoV antibody (Protein Data Bank [PDB] [71] ID: 6WS6 [65]). Antibody complementarity-determining (hypervariable) regions are critical in the recognition of antigens and extreme sequence variation in them allows antibodies to bind to a nearly limitless range of antigens. The particular antibody under study potently neutralises the SARS-CoV-2 virus [72]. The CDRH3 region is especially important for antigen recognition [73, 74], and its conformation is not restricted to a small set of canonical structures, unlike those of the other CDRs [73–75]. Note also that because Porter 5 is a machine learning-based algorithm, it is only expected to be accurate for sequences similar to natural sequences on which it was trained. Hence the need to use a naturally occurring protein, rather than a completely random sequence, for which Porter 5 may yield inaccurate SS predictions. The SS predicted by Porter 5 for the chosen CDRH3 is *x* =EEECCCCCCCCCCCCCCCCC, and this SS was highly accurate (85%) when compared to experimentally derived SS, EEECCCCCCCCCCCCCCEEE. We used site scanning [64] to generate a large number of different sequences within the neutral space of *x*. We then generated all mutant sequences reachable via a point mutation in DNA for a subset of the sample of neutral genotypes, and used Porter 5 to predict their SS (more details in SI I (D)).

Figure 2(a) shows the most frequent SS for each conditional complexity value found in the dataset generated via site scanning, as well as the estimated probability of transitioning from the starting phenotype *x* to all of these alternative ones. In Figure 2(b) we see a strong bias in transition probabilities with a linear decay in the upper bound to *P* (*x* → *y*), as predicted. The black line is a fit to the data (*a* = 1.6 and *b* = −2), added to highlight the upper bound. The panel also shows the predicted upper bound based on *a* = 1 and *b* = 0 (red line); this prediction is useful, but not as accurate as for RNA *L* = 40. Panel (c) presents the same data as Figure 2(a), except that 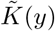 is on the horizontal axis instead of 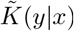. Interestingly, 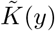 is quite similar to 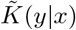, which is probably due to the fact that the original starting phenotype *x* is very simple (SI II). The scatter plot of 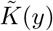 vs 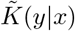 in Figure 2(d) shows that, as expected, there is a linear correlation between these two quantities (Pearson *r* = 0.87, p-value*<* 10^−6^). It is interesting that the correlation is stronger than for the RNA example, and this again can be rationalised by the fact that *x* is very simple. There do not appear to be any phenotypes for which 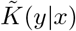 is much smaller than 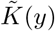. With probability-weighted sampling, the accuracy of predicting which of two phenotypes has higher probability is very high at 79%, and with uniform sampling is 77%. Figure 2(e) shows predictions for the ratio of the probabilities for transitioning to different phenotypes versus the difference between their estimated complexity in log scale. The latter displays a strong correlation (Pearson *r* = 0.93, p-value*<* 10^−6^). Finally, Figure 2(f) presents a histogram of conditional complexities, showing a stronger bias towards lower-complexity phenotypes than the data for RNA. In SI IV an additional supplementary figure shows accurate predictions for another protein SS example.

**FIG. 2.**
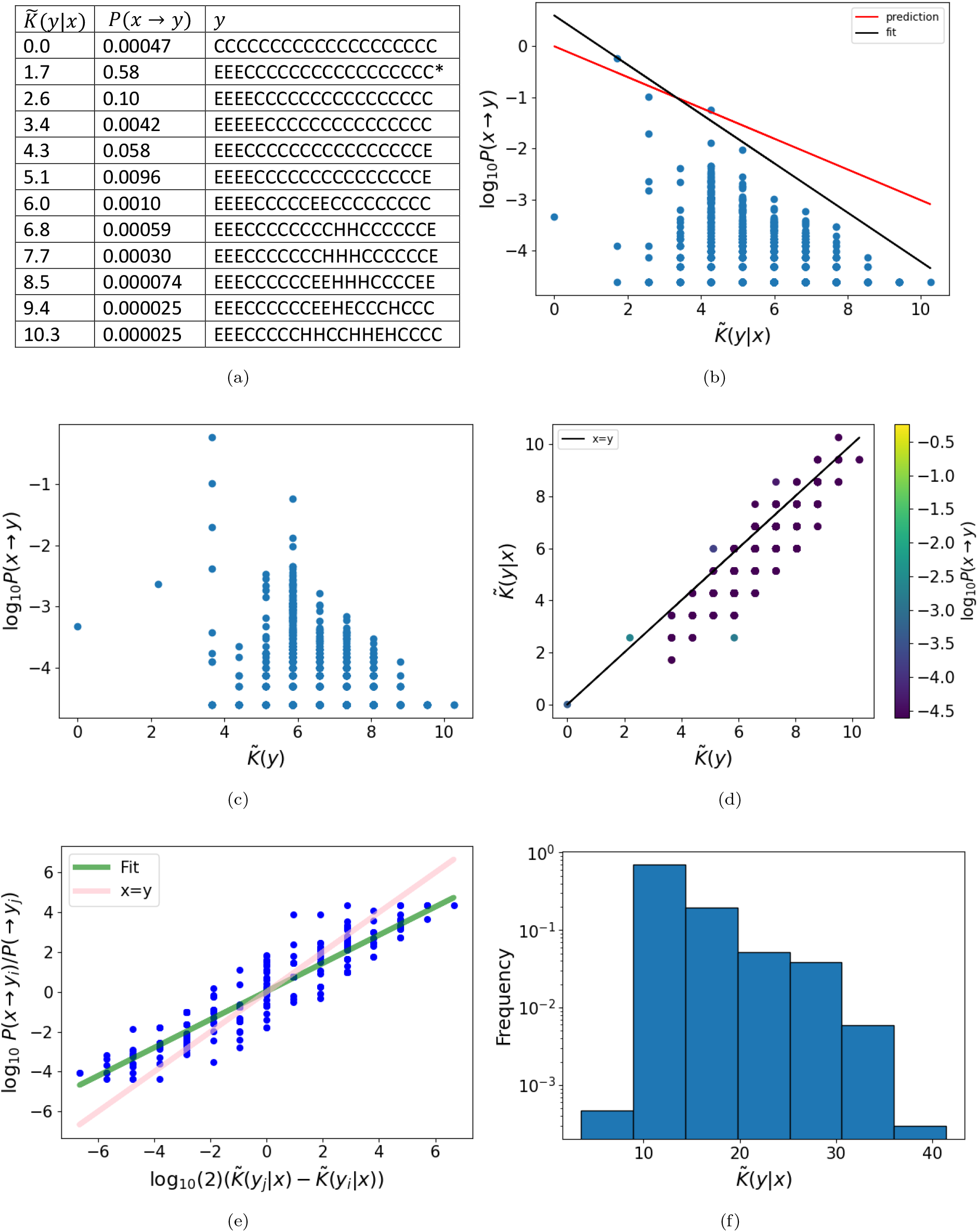
Protein secondary structure transition probabilities for CDRH3 from the heavy chain of a human monoclonal SARS-CoV antibody (PDB ID: 6WS6 [65]), length *L* = 20, *x* =EEECCCCCCCCCCCCCCCCC. (a) Table illustrating the highest-probability SS for each conditional complexity value. The starting phenotype *x* is marked with an asterisk (*), and *P* (*x* → *y*) is just the robustness since *x* = *y*. (b) Transition probabilities *P* (*x* → *y*) decrease exponentially with increasing conditional complexity 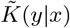. The phenotype with the lowest conditional complexity is CCCCCCCCCCCCCCCCCCCC. The phenotype with the highest probability is *y* = *x*. The predicted upper bound (red) is close to the fitted bound (black, cf. Eq. (10)). (c) The unconditional complexity 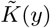 predicts the transition probabilities nearly as well as 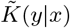, presumably because *x* has low complexity (see SI II). (d) 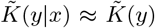 for most *y*, leading to a positive linear correlation between values. (e) Ratios of probabilities correlate strongly with differences in conditional complexity. (f) The histogram of conditional complexity values shows a bias towards simpler proteins with lower complexity values.

## VI. DISCUSSION

We have studied the problem of predicting the probability of transition between phenotypes *x* and *y, P* (*x* → *y*), from the perspective of algorithmic information theory (AIT), and specifically algorithmic probability estimates. We derived an upper bound on *P* (*x* → *y*) which depends on the conditional complexity of phenotype *y* given *x*. The derivations were motivated by the observation that the assignment of genotypes to phenotypes is highly structured, and the expectation that the constraints of information theory should therefore have predictive value in genotype-to-phenotype maps. Upon testing our various predictions on RNA and protein secondary structure examples, we found good quantitative agreement between the theory and the simulations.

The benefit of developing this theoretical approach is that it allows predictions to be made about transition probabilities for GP maps in which only phenotypes are observed and little or no knowledge about the map is available. This approach is also relevant to uncovering general common properties of GP maps, which are important for the advancement of the currently underdeveloped field of theoretical biology [76].

In this study we restricted our attention to one aspect of the structure of the GP, without including evolutionary dynamic effects resulting from mutation rates, strength of selection, population size or others that would be relevant in biology. Therefore we leave these to future work. It is interesting, however, that studies of natural RNA shapes [21, 22] and the shapes of protein complexes [23] have shown that GP map biases alone can be very good predictors of natural biological shape frequencies (see also refs. [41, 77–79] for more on different types of biases and evolutionary outcomes). Therefore, it may be that the transition probability biases discussed here, resulting from conditional complexity constraints, are strong enough that their stamp is still observable even in natural data. We suggest that an interesting follow-on study to ours would be to test this with natural bioinformatic databases.

In this study we have tested our transition probability predictions on two biologically important GP maps, namely RNA and protein sequence-to-structure maps. However, in these GP maps the connection between genotypes and phenotypes is quite direct and fairly simple. Furthermore, for computational reasons, we studied only short RNA and proteins. In biology, many GP maps have a less direct connection between genotypes and phenotypes, and it remains a possibility that our probability predictions do not work well for such complicated maps. We leave the exploration of the limits of applicability of our theory for future work. Having said this, it is noteworthy that other researchers have empirically observed a tendency for genetic mutations to favour simpler morphologies, specifically in teeth [80], embryo [81], and leaf shapes [82]. We believe that our results help rationalise these observations within a general theoretical framework. Additionally, these empirical observations made in the context of complicated and realistic biological maps may suggest that our theory can be applied in more complex GP maps.

Another area of potential applicability of our transition probability predictions is in genetic algorithms for optimisation. Indeed, Hu and Banzaff [83], as well as others [84, 85], have studied optimisation problems and shown that some target phenotypes are harder to find than others, not just because of having a low global frequency but also due to local mutational connections. In future work, it would be interesting to assess if these mutational connections are related to conditional complexity, as our theory would suggest.

Robustness to genetic mutations is an important property for organisms [8], but a general explanation for the high levels of robustness observed in GP maps has been lacking [50]. Our information theory perspective here relates to this question because we have seen that transitions to phenotypes which are similar to the starting phenotype tend to have high probability, and of course a phenotype is most similar to itself. We intend to explore this in detail in a forthcoming study.

Other authors have used information theory for non-UTMs to derive some results which are related to the ones we derived here. For example Calude et al [86] developed bounds for finite-state machines (the simplest computing devices), and moreover Merhav et al [87] have derived similar probability bounds to ours directly in terms of Lempel-Ziv complexity. It would be interesting to see if these calculations for non-UTMs could be extended to GP maps. In SI III (A) we discuss in more detail the use of AIT arguments in science.

While the upper bound from Eq. (10) appears to work well in the simulations presented here, a main weakness in our predictions is that many phenotypes that have low conditional complexities 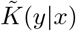 also have low probabilities. Because these phenotypes fall far below the upper bound, their precise probabilities are not well predicted by the theory. These low-complexity, low-probability patterns have been described as having low absolute information content, but due to map-specific biases they are ‘hard’ for the map to make, and hence have low probability [31]. The origins and nature of these types of patterns have been recently studied [62], but a full understanding of them and knowledge regarding how to improve probability predictions of these have not yet been achieved. Despite the challenge of low-complexity, low-probability patterns, we were still able to make high-accuracy upper bounds on phenotype probabilities as well as high-accuracy (*∼*80%) predictions about which of two phenotypes is more likely, just using complexity values.

The quantitatively accurate predictions we describe here motivate further investigation of the use of AIT-inspired predictions in biology, evolution, and other natural sciences.

## Acknowledgments and Funding Statement

This project has been partially supported by Gulf University for Science and Technology under project code: ISG — Case grant number 263301 and a Summer Faculty Fellowship (both awarded to KD). This work was performed using resources provided by the Cambridge Service for Data Driven Discovery (CSD3) operated by the University of Cambridge

Research Computing Service (www.csd3.cam.ac.uk), provided by Dell EMC and Intel using Tier-2 funding from the Engineering and Physical Sciences Research Council (capital grant EP/T022159/1), and DiRAC funding from the Science and Technology Facilities Council (www.dirac.ac.uk).

## Data availability

The data sets generated during and analysed during the current study are available from the corresponding author(s) on request.

## Author contributions

Conceived the study: KD, JN, SA, AL. Analytic calculations: KD. Simulations and data analysis: KD, JN. Wrote the paper: KD, JN, SA, AL.

## References

[1] Andreas Wagner. Arrival of the fittest: solving evolution’s greatest puzzle. Penguin, 2014.

[2] Sebastian E Ahnert. Structural properties of genotype– phenotype maps. Journal of The Royal Society Interface, 14(132):20170275, 2017.

[3] Susanna Manrubia, José A Cuesta, Jacobo Aguirre, Sebastian E Ahnert, Lee Altenberg, Alejandro V Cano, Pablo Catalán, Ramon Diaz-Uriarte, Santiago F Elena, Juan Antonio García-Martín, et al. From genotypes to organisms: State-of-the-art and perspectives of a cornerstone in evolutionary dynamics. Physics of Life Reviews, 38:55–106, 2021.

[4] P. Schuster, W. Fontana, P.F. Stadler, and I.L. Hofacker. From sequences to shapes and back: A case study in RNA secondary structures. Proceedings: Biological Sciences, 255(1344):279–284, 1994.

[5] E. Borenstein and D.C. Krakauer. An end to endless forms: epistasis, phenotype distribution bias, and nonuniform evolution. PLoS Comput Biol, 4(10):e1000202, 2008.

[6] I.G. Johnston, S.A. Ahnert, J.P.K. Doye, and A.A. Louis. Evolutionary dynamics in a simple model of self-assembly. Physical Review E, 83:066105, 2011.

[7] Kamaludin Dingle. Probabilistic bias in genotype-phenotype maps. PhD thesis, University of Oxford, 2014.

[8] A. Wagner. Robustness and evolvability in living systems. Princeton University Press Princeton, NJ:, 2005.

[9] T. Jorg, O.C. Martin, and A. Wagner. Neutral network sizes of biological RNA molecules can be computed and are not atypically small. BMC bioinformatics, 9(1):464, 2008.

[10] J.A. Draghi, T.L. Parsons, G.P. Wagner, and J.B. Plotkin. Mutational robustness can facilitate adaptation. Nature, 463(7279):353–355, 2010.

[11] Jacobo Aguirre, Javier M Buldú, Michael Stich, and Susanna C Manrubia. Topological structure of the space of phenotypes: the case of rna neutral networks. PloS one, 6(10):e26324, 2011.

[12] M.C. Cowperthwaite, E.P. Economo, W.R. Harcombe, E.L. Miller, and L.A. Meyers. The ascent of the abundant: how mutational networks constrain evolution. PLoS computational biology, 4(7):e1000110, 2008.

[13] Steffen Schaper and Ard A Louis. The arrival of the frequent: how bias in genotype-phenotype maps can steer populations to local optima. PloS one, 9(2):e86635, 2014.

[14] Sam F Greenbury, Iain G Johnston, Ard A Louis, and Sebastian E Ahnert. A tractable genotype–phenotype map modelling the self-assembly of protein quaternary structure. Journal of The Royal Society Interface, 11(95):20140249, 2014.

[15] Pablo Catalán, Susanna Manrubia, and José A Cuesta. Populations of genetic circuits are unable to find the fittest solution in a multilevel genotype–phenotype map. Journal of the Royal Society Interface, 17(167):20190843, 2020.

[16] S. Psujek and R.D. Beer. Developmental bias in evolution: evolutionary accessibility of phenotypes in a model evo-devo system. Evolution & Development, 10(3):375–390, 2008.

[17] Walter Fontana, Danielle AM Konings, Peter F Stadler, and Peter Schuster. Statistics of rna secondary structures. Biopolymers: Original Research on Biomolecules, 33(9):1389–1404, 1993.

[18] Erik A Schultes, Alexander Spasic, Udayan Mohanty, and David P Bartel. Compact and ordered collapse of randomly generated rna sequences. Nature structural & molecular biology, 12(12):1130–1136, 2005.

[19] Sandra Smit, Michael Yarus, and Rob Knight. Natural selection is not required to explain universal compositional patterns in rrna secondary structure categories. RNA, 12(1):1–14, 2006.

[20] Kamaludin Dingle, Steffen Schaper, and Ard A Louis. The structure of the genotype–phenotype map strongly constrains the evolution of non-coding RNA. Interface focus, 5(6):20150053, 2015.

[21] Kamaludin Dingle, Fatme Ghaddar, Petr Šulc, and Ard A Louis. Phenotype bias determines how natural rna structures occupy the morphospace of all possible shapes. Molecular biology and evolution, 39(1):msab280, 2022.

[22] Fatme Ghaddar and Kamaludin Dingle. Random and natural non-coding rna have similar structural motif patterns but can be distinguished by bulge, loop, and bond counts. bioRxiv, 2022.

[23] Iain G Johnston, Kamaludin Dingle, Sam F Greenbury, Chico Q Camargo, Jonathan PK Doye, Sebastian E Ahnert, and Ard A Louis. Symmetry and simplicity spontaneously emerge from the algorithmic nature of evolution. Proceedings of the National Academy of Sciences, 119(11):e2113883119, 2022.

[24] Jasper Franke, Alexander Klözer, J Arjan GM de Visser, and Joachim Krug. Evolutionary accessibility of mutational pathways. PLoS Comput Biol, 7(8):e1002134, 2011.

[25] L. Tan, S. Serene, H.X. Chao, and J. Gore. Hidden randomness between fitness landscapes limits reverse evolution. Physical Review Letters, 106(19):198102, 2011.

[26] Isaac Salazar-Ciudad and Miquel Marín-Riera. Adaptive dy-namics under development-based genotype–phenotype maps. Nature, 497(7449):361–364, 2013.

[27] R. J. Solomonoff. A preliminary report on a general theory of inductive inference (revision of report v-131). Contract AF, 49(639):376, 1960.

[28] A.N. Kolmogorov. Three approaches to the quantitative definition of information. Problems of information transmission, 1(1):1–7, 1965.

[29] Gregory J Chaitin. A theory of program size formally identical to information theory. Journal of the ACM (JACM), 22(3):329–340, 1975.

[30] Kamaludin Dingle, Chico Q Camargo, and Ard A Louis. Input–output maps are strongly biased towards simple outputs. Nature communications, 9(1):761, 2018.

[31] Kamaludin Dingle, Guillermo Valle Pérez, and Ard A Louis. Generic predictions of output probability based on complexities of inputs and outputs. Scientific reports, 10(1):1–9, 2020.

[32] SE Ahnert, IG Johnston, TMA Fink, JPK Doye, and AA Louis. Self-assembly, modularity, and physical complexity. Physical Review E, 82(2):026117, 2010.

[33] C.H. Bennett. The thermodynamics of computation – a review. International Journal of Theoretical Physics, 21(12):905–940, 1982.

[34] Artemy Kolchinsky and David H Wolpert. Thermodynamic costs of turing machines. Physical Review Research, 2(3):033312, 2020.

[35] W.H. Zurek. Algorithmic randomness and physical entropy. Physical Review A, 40(8):4731, 1989.

[36] Markus P Mueller. Law without law: from observer states to physics via algorithmic information theory. Quantum, 4:301, 2020.

[37] Ram Avinery, Micha Kornreich, and Roy Beck. Universal and accessible entropy estimation using a compression algorithm. Physical review letters, 123(17):178102, 2019.

[38] Stefano Martiniani, Paul M Chaikin, and Dov Levine. Quantifying hidden order out of equilibrium. Physical Review X, 9(1):011031, 2019.

[39] P. Ferragina, R. Giancarlo, V. Greco, G. Manzini, and G. Valiente. Compression-based classification of biological sequences and structures via the universal similarity metric: experimental assessment. BMC bioinformatics, 8(1):252, 2007.

[40] Alyssa Adams, Hector Zenil, Paul CW Davies, and Sara Imari Walker. Formal definitions of unbounded evolution and innovation reveal universal mechanisms for openended evolution in dynamical systems. Scientific reports, 7(1):1–15, 2017.

[41] Kamaludin Dingle. Fitness, optima, and simplicity. Preprints, 2022080402, 2022.

[42] Hector Zenil, Fernando Soler-Toscano, Kamaludin Dingle, and Ard A Louis. Correlation of automorphism group size and topological properties with program-size complexity evaluations of graphs and complex networks. Physica A: Statistical Mechanics and its Applications, 404:341–358, 2014.

[43] Hector Zenil, Narsis A Kiani, and Jesper Tegnér. A review of graph and network complexity from an algorithmic information perspective. Entropy, 20(8):551, 2018.

[44] Paul MB Vitányi. Similarity and denoising. Philosophical Transactions of the Royal Society A: Mathematical, Physical and Engineering Sciences, 371(1984), 2013.

[45] R. Cilibrasi and P.M.B. Vitányi. Clustering by compression. Information Theory, IEEE Transactions on, 51(4):1523–1545, 2005.

[46] Hector Zenil, Narsis A Kiani, Allan A Zea, and Jesper Tegnér. Causal deconvolution by algorithmic generative models. Nature Machine Intelligence, 1(1):58–66, 2019.

[47] H. Zenil and J.P. Delahaye. An algorithmic information theoretic approach to the behaviour of financial markets. Journal of Economic Surveys, 25(3):431–463, 2011.

[48] Kamaludin Dingle, Rafiq Kamal, and Boumediene Hamzi. A note on a priori forecasting and simplicity bias in time series. arXiv preprint arXiv:2203.05391, 2022.

[49] M. Li and P.M.B. Vitanyi. An introduction to Kolmogorov complexity and its applications. Springer-Verlag New York Inc, 2008.

[50] Sam F Greenbury, Steffen Schaper, Sebastian E Ahnert, and Ard A Louis. Genetic correlations greatly increase mutational robustness and can both reduce and enhance evolvability. PLoS computational biology, 12(3):e1004773, 2016.

[51] Marcel Weiß and Sebastian E Ahnert. Phenotypes can be robust and evolvable if mutations have non-local effects on sequence constraints. Journal of The Royal Society Interface, 15(138):20170618, 2018.

[52] Alan Mathison Turing. On computable numbers, with an application to the entscheidungsproblem. J. of Math, 58(345-363):5, 1936.

[53] C.S. Calude. Information and randomness: An algorithmic perspective. Springer, 2002.

[54] P. Gács. Lecture notes on descriptional complexity and randomness. Boston University, Graduate School of Arts and Sciences, Computer Science Department, 1988.

[55] Alexander Shen, Vladimir A Uspensky, and Nikolay Vereshchagin. Kolmogorov complexity and algorithmic randomness, volume 220. American Mathematical Society, 2022.

[56] Sean D Devine. Algorithmic Information Theory for Physicists and Natural Scientists. IOP Publishing, 2020.

[57] L.A. Levin. Laws of information conservation (nongrowth) and aspects of the foundation of probability theory. Problemy Peredachi Informatsii, 10(3):30–35, 1974.

[58] Ray J Solomonoff. The discovery of algorithmic probability. Journal of Computer and System Sciences, 55(1):73–88, 1997.

[59] Paul MB Vitányi. Conditional kolmogorov complexity and universal probability. Theoretical Computer Science, 501:93–100, 2013.

[60] A. Lempel and J. Ziv. On the complexity of finite sequences. Information Theory, IEEE Transactions on, 22(1):75–81, 1976.

[61] Ming Li, Xin Chen, Xin Li, Bin Ma, and Paul MB Vitányi. The similarity metric. IEEE transactions on Information Theory, 50(12):3250–3264, 2004.

[62] Mohamed Alaskandarani and Kamaludin Dingle. Low complexity, low probability patterns and consequences for algorithmic probability applications. arXiv preprint arXiv:2207.12251, 2022.

[63] Ronny Lorenz, Stephan H Bernhart, Christian Höner Zu Siederdissen, Hakim Tafer, Christoph Flamm, Peter F Stadler, and Ivo L Hofacker. Viennarna package 2.0. Algorithms for molecular biology, 6(1):26, 2011.

[64] Marcel Weiß and Sebastian E Ahnert. Using small samples to estimate neutral component size and robustness in the genotype–phenotype map of rna secondary structure. Journal of the Royal Society Interface, 17(166):20190784, 2020.

[65] D. Pinto, D., Park, Y.J., Beltramello, M., Walls, A.C., Tortorici, M.A., Bianchi, S., Jaconi, S., Culap, K., Zatta, F., Marco, A.D., Peter, A., Guarino, B., Spreafico, R., Cameroni, E., Case, J.B., Chen, R.E., Havenar-Daughton, C., Snell, G., Telenti, A. Vi. Structural and functional analysis of a potent sarbecovirus neutralizing antibody, DOI: 10.2210/pdb6WS6/pdb, 2020.

[66] Cyrus Chothia. One thousand families for the molecular biologist. Nature, 357(6379):543–544, 1992.

[67] Yong Yun Ji and You Quan Li. The role of secondary structure in protein structure selection. Eur. Phys. J. E, 32(1):103–107, 2010.

[68] John Jumper, Richard Evans, Alexander Pritzel, Tim Green, Michael Figurnov, Olaf Ronneberger, Kathryn Tunyasuvunakool, Russ Bates, Augustin Žídek, Anna Potapenko, et al. Highly accurate protein structure prediction with alphafold. Nature, 596(7873):583–589, 2021.

[69] Mirko Torrisi, Manaz Kaleel, and Gianluca Pollastri. Deeper profiles and cascaded recurrent and convolutional neural networks for state-of-the-art protein secondary structure prediction. Scientific reports, 9(1):1–12, 2019.

[70] Gwen R. Buel and Kylie J. Walters. Can AlphaFold2 predict the impact of missense mutations on structure? Nat. Struct. Mol. Biol., 29(1):1–2, jan 2022.

[71] Helen M Berman, John Westbrook, Zukang Feng, Gary Gilliland, Talapady N Bhat, Helge Weissig, Ilya N Shindyalov, and Philip E Bourne. The protein data bank. Nucleic acids research, 28(1):235–242, 2000.

[72] Dora Pinto, Young Jun Park, Martina Beltramello, Alexandra C. Walls, M. Alejandra Tortorici, Siro Bianchi, Stefano Jaconi, Katja Culap, Fabrizia Zatta, Anna De Marco, Alessia Peter, Barbara Guarino, Roberto Spreafico, Elisabetta Cameroni, James Brett Case, Rita E. Chen, Colin Havenar-Daughton, Gyorgy Snell, Amalio Telenti, Herbert W. Virgin, Antonio Lanzavecchia, Michael S. Diamond, Katja Fink, David Veesler, and Davide Corti. Cross-neutralization of SARS-CoV-2 by a human monoclonal SARS-CoV antibody. Nature, 583(7815):290–295, 2020.

[73] Ian A. Wilson and Robyn L Stanfield. Antibody-antigen interactions: new structures and new conformational changes. Curr. Opin. Struct. Biol., 4(6):857–867, jan 1994.

[74] Brian D. Weitzner, Roland L. Dunbrack, and Jeffrey J. Gray. The Origin of CDR H3 Structural Diversity. Structure, 23(2):302–311, feb 2015.

[75] R. J. Pantazes and C. D. Maranas. OptCDR: A general computational method for the design of antibody complementarity determining regions for targeted epitope binding. Protein Eng. Des. Sel., 23(11):849–858, 2010.

[76] David C Krakauer, James P Collins, Douglas Erwin, Jessica C Flack, Walter Fontana, Manfred D Laubichler, Sonja J Prohaska, Geoffrey B West, and Peter F Stadler. The challenges and scope of theoretical biology. Journal of theoretical biology, 276(1):269–276, 2011.

[77] L.Y. Yampolsky and A. Stoltzfus. Bias in the introduction of variation as an orienting factor in evolution. Evolution & Development, 3(2):73–83, 2001.

[78] Alejandro V Cano, Hana Rozhoňová, Arlin Stoltzfus, David M McCandlish, and Joshua L Payne. Mutation bias shapes the spectrum of adaptive substitutions. Proceedings of the National Academy of Sciences, 119(7):e2119720119, 2022.

[79] Isaac Salazar-Ciudad. Why call it developmental bias when it is just development? Biology Direct, 16:1–13, 2021.

[80] Enni Harjunmaa, Aki Kallonen, Maria Voutilainen, Keijo Hämäläinen, Marja L. Mikkola, and Jukka Jernvall. On the difficulty of increasing dental complexity. Nature, 483(7389):324–327, 2012.

[81] Pascal F Hagolani, Roland Zimm, Renske Vroomans, and Isaac Salazar-Ciudad. On the evolution and development of morphological complexity: A view from gene regulatory networks. PLoS computational biology, 17(2):e1008570, 2021.

[82] R. Geeta, L.M. Davalos, A. Levy, L. Bohs, M. Lavin, K. Mummenhoff, N. Sinha, and M.F. Wojciechowski. Keeping it simple: flowering plants tend to retain, and revert to, simple leaves. New Phytologist, 2011.

[83] Ting Hu, Marco Tomassini, and Wolfgang Banzhaf. A network perspective on genotype–phenotype mapping in genetic programming. Genetic Programming and Evolvable Machines, 21(3):375–397, 2020.

[84] Wolfgang Banzhaf. Genotype-phenotype-mapping and neutral variation—a case study in genetic programming. In International Conference on Parallel Problem Solving from Nature, pages 322–332. Springer, 1994.

[85] Peter A Whigham, Grant Dick, and James Maclaurin. On the mapping of genotype to phenotype in evolutionary algorithms. Genetic Programming and Evolvable Machines, 18(3):353–361, 2017.

[86] Cristian S Calude, Kai Salomaa, and Tania K Roblot. Finite state complexity. Theoretical Computer Science, 412(41):5668–5677, 2011.

[87] Neri Merhav and Asaf Cohen. Universal randomized guessing with application to asynchronous decentralized brute– force attacks. IEEE Transactions on Information Theory, 66(1):114–129, 2019.

