## Supplementary text and figures for "Predicting phenotype transition probabilities via conditional algorithmic probability approximations"

### Supplementary Information for: Predicting phenotype transition probabilities via conditional algorithmic probability approximations —Published in *Journal of the Royal Society Interface*—

Kamaludin Dingle,[1,2,3] Javor K. Novev,[1] Sebastian E. Ahnert,[1] and Ard A. Louis[4]

[1] *Department of Chemical Engineering and Biotechnology, Cambridge University, UK*
[2] *Department of Computing and Mathematical Sciences, California Institute of Technology, USA*
[3] *Centre for Applied Mathematics and Bioinformatics (CAMB), Department of Mathematics and Natural Sciences, Gulf University for Science and Technology, Kuwait*
[4] *Rudolf Peierls Centre for Theoretical Physics, Department of Physics, Oxford University, UK*

(Dated: November 13, 2022)

#### I. METHODS

##### A. Complexity

To approximate the Kolmogorov complexity we followed the methodology of ref. [1], in which the function

$$C_{LZ}(x) = \begin{cases} \log_2(n), & x = 0^n \text{ or } 1^n \\ \log_2(n)[N_w(x_1...x_n) + N_w(x_n...x_1)]/2, & \text{otherwise} \end{cases} \tag{S.1}$$

was defined where $N_w$ forms the basis for the 1976 Lempel and Ziv complexity measure [2]. Here the simplest strings $0^n$ and $1^n$ are separated out because $N_w(x)$ assigns complexity $K = 1$ to the string 0 or 1, but complexity 2 to $0^n$ or $1^n$ for $n \geq 2$, whereas the true Kolmogorov complexity of such a trivial string actually scales as $\log_2(n)$ for typical $n$, because one only needs to encode $n$. The minimum possible value is $K(x){\approx}0$ for a simple set, and so, e.g. for binary strings of length $n$ we can expect $0 \lesssim K(x) \lesssim n$ bits. Because for a random string of length $n$ the value $C_{LZ}(x)$ is often much larger than $n$, especially for short strings, we scale the complexity

$$\tilde{K}(x) = \log_2(M) \cdot \frac{C_{LZ}(x) - \min_x(C_{LZ})}{\max_x(C_{LZ}) - \min_x(C_{LZ})} \tag{S.2}$$

where $M$ is the maximum possible number of phenotypes in the system, and the minimum and maximum complexities are over all strings $x$ which the map can generate. $\tilde{K}(x)$ is the approximation to Kolmogorov complexity that we use throughout. This scaling results in $0 \leq \tilde{K}(x) \leq n$ which is the desirable range of values.

##### B. Predicting which of two transitions is more likely

Another quantity that may be predicted using the theory from Section IV in the main text is the ratio of probabilities for transitioning from one phenotype to different alternative phenotypes. In this section, we describe a method for such predictions, which we test numerically in the main text.

Call $y_i$ the resulting phenotype after a single point mutation to a randomly chosen genotype in the neutral set (NS) of $x$. Call $y_j$ the resulting phenotype after a single point mutation to another independently chosen random genotype, also in the NS of $x$. We can use the theory from Section IV in the paper to predict which of the two phenotypes $y_i$ and $y_j$ has a higher probability directly from complexity estimates. This is interesting because it is often valuable to know whether $P(x \to y_i) > P(x \to y_j)$ or $P(x \to y_i) < P(x \to y_j)$, rather than trying to guess the exact values of $P(x \to y_i)$ and $P(x \to y_j)$. Fortunately, the constants $a$ and $b$ are not required only the relative value of $\tilde{K}(y_i|x)$ and $\tilde{K}(y_j|x)$ determines whether $2^{-a\tilde{K}(y_i|x)-b}$ or $2^{-a\tilde{K}(y_j|x)-b}$ is larger. So even if we could not estimate $a$ or $b$ accurately, we could still make a prediction about which phenotype is more likely to arise through a point mutation of $x$

The approach we use is to predict:

$$\begin{aligned} P(x \to y_i) > P(x \to y_j) &\quad \texttt{if} \quad \tilde{K}(y_i|x) < \tilde{K}(y_j|x) \\ P(x \to y_i) < P(x \to y_j) &\quad \texttt{if} \quad \tilde{K}(y_i|x) > \tilde{K}(y_j|x) \\ P(x \to y_i) \geq P(x \to y_j) &\quad \texttt{w/prob 0.5 if} \quad \tilde{K}(y_i|x) = \tilde{K}(y_j|x) \end{aligned}$$

For the last condition, $\tilde{K}(y_i|x) = \tilde{K}(y_j|x)$, a random number (e.g., coin flip) is chosen and used to predict whether $P(x \to y_i) \geq P(x \to y_j)$ or not, each with 50% probability. It may be suggested that predicting $P(x \to y_i) = P(x \to y_j)$ is more appropriate if the two conditional complexities are equal, but following this suggestion will lead to many erroneous predictions. This is because there are very many unique probability values, and so it is highly unlikely that $P(x \to y_i) = P(x \to y_j)$ exactly, whereas the complexities are far more coarsely measured, and hence coincidences of complexity values are much more likely. According to this prediction protocol, if complexity had no role in modulating probabilities, then the null success rate of correctly guessing which of $P(x \to y_i)$ or $P(x \to y_j)$ is higher should be roughly 50% (if all probabilities were distinct it would be 50% exactly, but some of the probabilities are repeated).

##### C. RNA

We used a (pseudo) random number generator to generate nucleotide sequences of length $L = 40$. We predicted the RNA secondary structures (SS) with the RNA Vienna folding package [3], which uses a thermodynamics-based algorithm, downloaded from `https://anaconda.org/bioconda/viennarna`. In order to estimate transition probabilities $P(x \to y)$, a sample of sequences from the neutral space (NS) of the phenotype $x$ are required. Generating this sample by storing random sequences and saving only those which fold to phenotype $x$ would be a possibility in principle, but it is highly computationally taxing. Hence we used site scanning [4], a method for sampling from the NS of a given phenotype. The site-scanning method introduces a random point mutation in the first site of the initial genotype; if this mutation is not neutral, it is reverted and a different random mutation is introduced at the same site. This process is repeated until a neutral mutant is identified or all possible point mutations at the first site have been exhausted, whereupon the algorithm moves on to the next site; when the algorithm reaches the last site, it restarts from the first one, and the process continues until the desired number of neutral mutants has been derived.

For the RNA figure in the main text, we used $10^7$ 'steps' in the site-scanning algorithm, and saved only a randomly chosen fraction of 0.1 of the sequences - a subsample in the terminology of [4] - in order to ensure low correlations between sequences in the sample. For the RNA figure in the appendix, we used $10^6$ 'steps'. For each of the saved sequences in the sample, we introduced a single random point mutation, and used the resulting mutated sequences to predict the resulting RNA SS. The frequency of each SS was used to define $P(x \to y)$.

##### D. Proteins

We predicted protein SS with the fast version of the machine learning-based tool Porter 5, which makes use of protein alignments performed via HHblits [5]. For each studied protein, we used the site-scanning technique as described in the preceding section with minor modifications as follows. 1) At each site in a given protein sequence, we introduced point mutations that are accessible via a point mutation in the DNA sequence following the standard genetic code. In doing so, we considered all such mutations equally likely without weighting for the number of codons that code for a particular amino acid or for codon bias; moreover, we disregarded mutations that change a codon encoding an amino acid to a stop codon. 2) Instead of modifying the entire protein, we only looked at mutations in the H3 hypervariable region of the heavy chains of two antibody proteins, and if the algorithm reached a dead end (i.e., a sequence with no neutral point mutants), we restarted it from a random genotype that had been derived earlier in the run. 3) When generating neutral genotypes during the site-scanning walk, we discarded genotypes that had previously been encountered in the run, instead considering ones that feature alternative amino acid residues at the site under consideration or, if all such residues have been exhausted, moving on to the next site.

We generated a sample of 4000 neutral genotypes, from which we then generated the complete point-mutational neighborhood of a random subsample of 200 sequences, corresponding to a fraction of 0.05 of the entire sample. Figure 2 in the main text is based on the predictions of the SS of all 40402 unique sequences generated during the site-scanning run - this includes the 4000 neutral mutants in the sample, the non-neutral ones encountered while generating the sample, and the mutants accessible through point DNA mutations from the neutral ones in the subsample. We use the CDRH3 of the human SARS-CoV antibody with PDB entry 6WS6 [6, 7] as the starting point for the run. The SS predictions for the CDRH3 are from running the Porter 5 algorithm on the full sequence of the antibody's heavy chain. For the wild-type H3 region of 6WS6, this results in an accuracy of 85% when compared with the experimental SS data in PDB. The primary structure for the region under study (using single-letter amino acid codes) is `ARDYTRGAWFGESLIGGFDN` [7], the secondary structure as recorded in the PDB entry - `EEECCCCCCCCCCCCCCEEE`, and the fast Porter 5 prediction based on the full sequence of the heavy chain - `EEECCCCCCCCCCCCCCCCC`.

For the CDRH3 of HIV-1-neutralizing antibody PG16 (PDB entry: 4DQO [8, 9]), we applied the same approach, except that we used the fast Porter 5 predictions for just the H3 region, which was much less computationally intensive but resulted in only 36% accuracy in predicting the SS of the wild type. The primary structure of the wild-type CDRH3 is `AREAGGPIWHDDVKYYDFNDGYYNYHYMDV` according to the IMGT definition as recorded in the structural antibody database SAbDab [10], the SS predicted by Porter 5 is `CCCCCCCCCCCCCEEEECCCCEEEEEECCC`, and the SS from PDB is `EEEEEECEEECCEEEEHHHHCEEEEEEE` [8]. The site-scanning run and the subsequent exploration of the point-mutational neighborhoods of neutral genotypes from the subsample generated 107330 mutant sequences. The data for this protein is presented in the section below on Supplementary Figures.

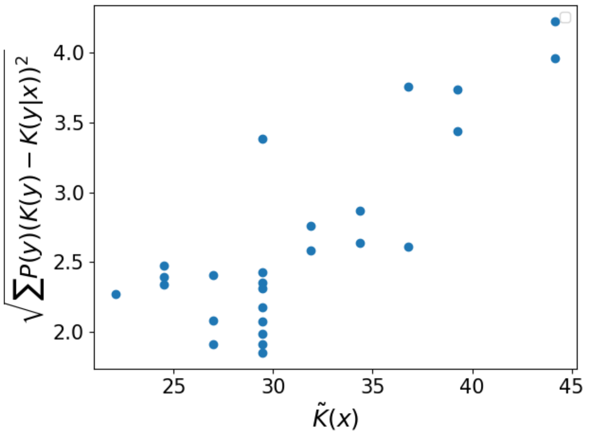


(a)

FIG. II.1. **The conditional complexity $\tilde{K}(y|x)$ is typically more different to $\tilde{K}(y)$ for more complex starting phenotypes $x$.** Using RNA $L = 30$ SS, we measure the average difference between $\tilde{K}(y|x)$ and $\tilde{K}(y)$. As expected from the discussion around Eq. (S.8), for simpler $x$ the difference is typically small, while for more complex $x$ the difference increases. The linear correlation has coefficient $r$= 0.81 with $p$-value$< 10^{-6}$.

#### II. EFFECT OF THE COMPLEXITY OF THE STARTING PHENOTYPE $x$

In the discussion around SI Eq. (S.8), we suggested that for a very simple starting phenotype $x$, the conditional and unconditional complexities will likely be similar, i.e. $\tilde{K}(y|x) \approx \tilde{K}(y)$. For complex $x$, we suggested that potentially $\tilde{K}(y|x) \ll \tilde{K}(y)$ for some $y$, especially those that are similar to $x$. Even though $\tilde{K}(y|x)$ and $\tilde{K}(y)$ may differ substantially only for a small fraction of possible $y$, because of the exponential dependence of $P(x \to y)$ on the value of $\tilde{K}(y|x)$, the small fraction of SS may potentially account for a large fraction of the probability $P(x \to y)$. As a brief investigation into the effect of the complexity of $x$ on the similarity of $\tilde{K}(y|x)$ and $\tilde{K}(y)$, in SI Figure II.1 we show a plot of the root-mean-square difference between these values, i.e. $\sqrt{\sum_y P(y)(\tilde{K}(y) - \tilde{K}(y|x))^2}$, vs the complexity of $x$. The Figure shows clearly that for lower complexity $x$, the difference is small, as expected. Interestingly, the difference grows linearly with increasing complexity of $x$ (linear correlation $r$= 0.81 with $p$-value$< 10^{-6}$). To generate these data, we used 25 RNA SS of length $L = 30$, with 20 000 steps in the site-scanning method saving a fraction of 0.1 of the sequences, i.e. the same methodology as described above in Section I C.

#### III. THEORY DETAILS

##### A. Can we use AIT in the natural sciences?

AIT is an abstract theory developed within theoretical computer science. It is far from obvious that AIT should be applicable in real-world settings, like biological GP maps. Indeed, the application of AIT to real-world science problems suffers from several problems, including that (a) Kolmogorov complexity is uncomputable, (b) the results are framed and proved in the context of universal Turing machine (UTMs) while many real world maps are not Turing complete, and (c) results are valid up to $O(1)$ terms and therefore, strictly, only accurate in the asymptotic limit of large complexities. Given these, it is surprising that AIT can be successfully applied at all.

The logic of the present study and Refs. [1, 11, 12] is that we take AIT as a theoretical framework to make predictions and then test those predictions in practical settings, while being aware of the fact that, strictly, the predictions have not been proven to hold in the regime in which we apply them. We know that for asymptotically large patterns, the results from AIT apply, because we can ignore $O(1)$ terms in the asymptotic regime. Despite not being in the asymptotic regime in, e.g., biological GP map studies, we use AIT results and empirically observe that they seem to work quite well, in the sense of making non-trivial predictions.

It is not entirely clear to us why these AIT predictions work so well, and why we can 'ignore' these $O(1)$ terms, which could in principle dominate. However, ignoring these $O(1)$ terms is not specific to our work: in much of computer science, as well as often in applied mathematics, and also in physics, theory is developed or proven in asymptotic regimes and then applied in practical, non-asymptotic, settings. In fact it is quite common to see results that are proved in the infinite limit, but work quite well far outside that regime. Why this is possible is an open question. Computer scientist Scott Aaronson [13] has pointed out that it is something of a mystery why merely examining the scaling form of equations is often good

| $\tilde{K}(y\|x)$ | $P(x \rightarrow y)$ | $y$ |
|---|---|---|
| 0.0 | 0.055 | ........................................ |
| 0.8 | 0.34 | .((.....(((((.......)))))........))..... * |
| 1.7 | 0.018 | (((.....(((((.......)))))........))).... |
| 2.6 | 0.0056 | .......((((((.......)))))).............. |
| 3.4 | 0.0012 | .....((.(((((.......))))))))............. |
| 4.3 | 0.015 | .((.....((((((.....))))))........))..... |
| 5.2 | 0.0080 | .((.....((((.........))))........))..... |
| 6.0 | 0.0016 | .((((...(((((.......))))).....)).))..... |
| 6.9 | 0.0033 | .((...(((((((.......)))))))......))..... |
| 7.8 | 0.0023 | .((..((.(((((.......))))).)).....))..... |
| 8.6 | 0.00088 | .((.(((.(((((.......)))))..)))...))..... |
| 9.5 | 0.00017 | .((..(((((((.......))))))......)))..)).. |
| 10.3 | 0.000050 | .((......(.(.(((....))).).)......))..... |
| 11.2 | 0.000090 | .((....(((.(.(........).).)))....))..... |
| 12.1 | 0.000010 | (((((.(..(((.....((......)).)))..))).))) |
| 12.9 | 0.000010 | ((((..((((.....)).))))))((.(((...))).)). |
| 13.8 | 0.000010 | (((.(.....).))).(((.(((..(....)..)))))). |
| 14.7 | 0.000010 | ((.((...)).)).(((.(((..(......)..))).))) |

(a)

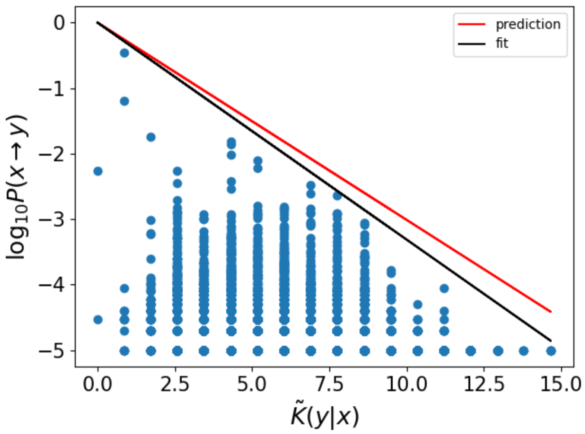


(b)

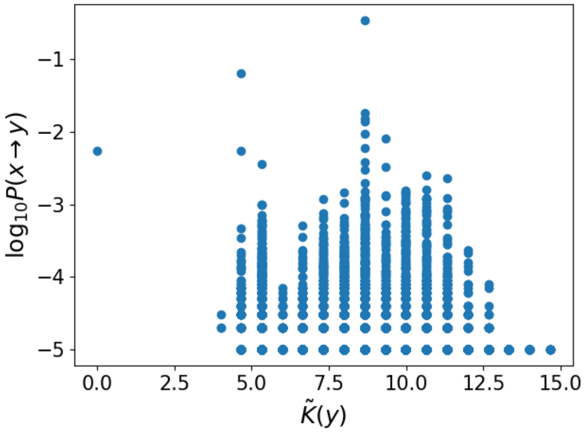


(c)

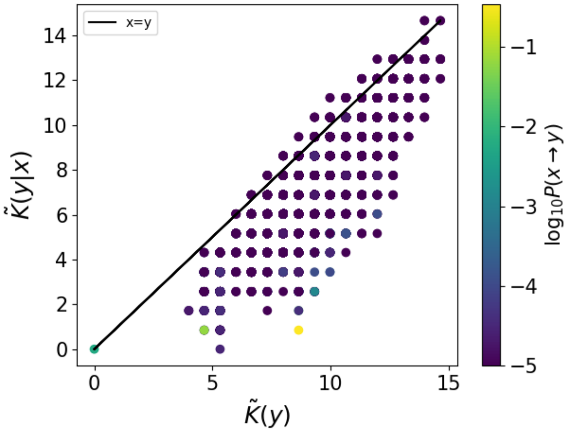


(d)

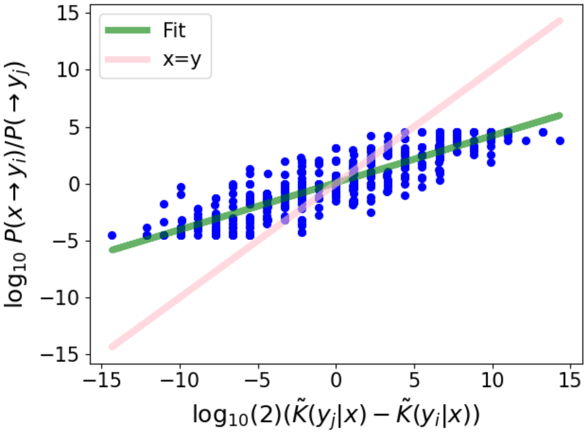


(e)

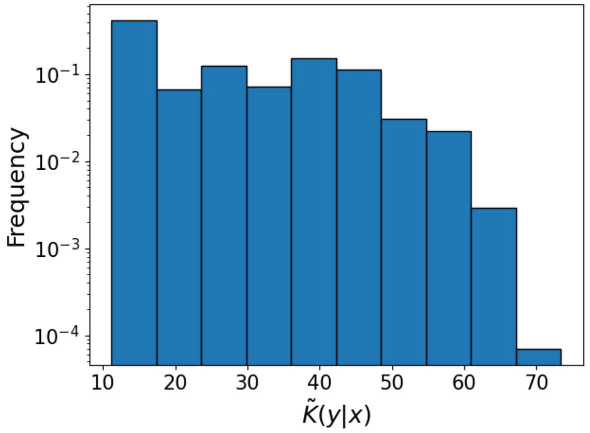


(f)

FIG. II.2. **RNA secondary structure transition probabilities for a sequence of length $L = 40$ nucleotides.** $x =$ `.((.....(((((.......)))))........)).....` (a) Table illustrating the SS with the highest probability for each complexity value. The starting phenotype $x$ is marked with an asterisk (*). (b) Transition probabilities $P(x \rightarrow y)$ decrease exponentially with increasing conditional complexity $\tilde{K}(y|x)$, upper bound depicted in black. The predicted upper bound (red) and the fitted bound (black) are close. (c) The unconditional complexity $\tilde{K}(y)$ does not predict the transition probabilities well. (d) $\tilde{K}(y|x) \approx \tilde{K}(y)$ for most $y$, leading to a positive linear correlation between values. (e) Ratios of probabilities correlate strongly with differences in conditional complexity. (f) The histogram of conditional complexity values shows a roughly flat distribution (on a log scale), but with some bias towards simplicity.

| $\tilde{K}(y\|x)$ | $P(x \to y)$ | $y$ |
|---|---|---|
| 0.0 | 0.0043 | CCCCCCCCCCCCCCCCCCCCCCCCCCCCCC |
| 1.0 | 0.38 | CCCCCCCCCCCCCEEEECCCCEEEEEECCC* |
| 1.9 | 0.021 | CCCCCCCCCCCCCEEEECCCCEEEEECCCC |
| 2.9 | 0.042 | CCCCCCCCCCCCCCEEECCCCEEEEEECCC |
| 3.8 | 0.027 | CCCCCCCCCCCCCEEEECCCCCEEEEECCC |
| 4.8 | 0.014 | CCCCCCCCCCCCCCEEECCCCCEEEEECCC |
| 5.8 | 0.0051 | CCCCCCCCCCCCCCEECCCCCCEEEEECCC |
| 6.7 | 0.00073 | CCCCCCCCCCCCCEEEECCCCEEEHEECCC |
| 7.7 | 0.00025 | CCCCCCCCCCCCCCEEECCCCEEEHEECCC |
| 8.7 | 0.000093 | CCCCCCCHHHHHHHHEECCCCCEEEEECCC |
| 9.6 | 0.000047 | CCCCCCCCCCCCCCEECCCCCEEEHEHCCC |
| 10.6 | 0.000019 | CCCCCCCCCCCCCEEEHHCCCEEHHEHCCC |
| 11.5 | 0.0000093 | CCCCCCHHHHCCCCHEHCCCCCEEEHECCC |
| 12.5 | 0.0000093 | CCCCCHHHCCCCCCEHHHCCHHHHEEHCCC |

(a)

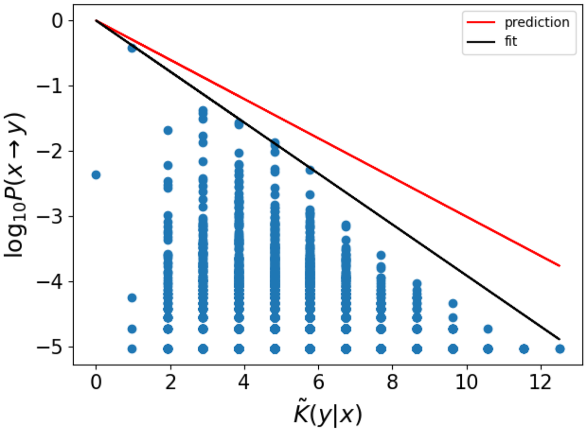


(b)

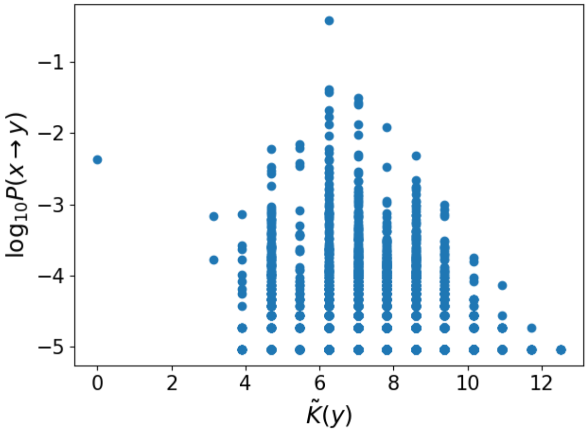


(c)

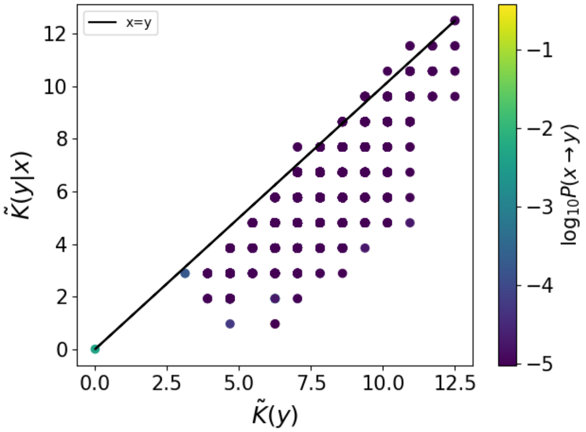


(d)

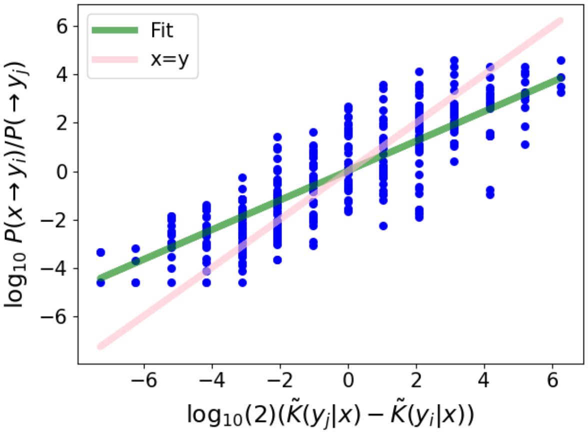


(e)

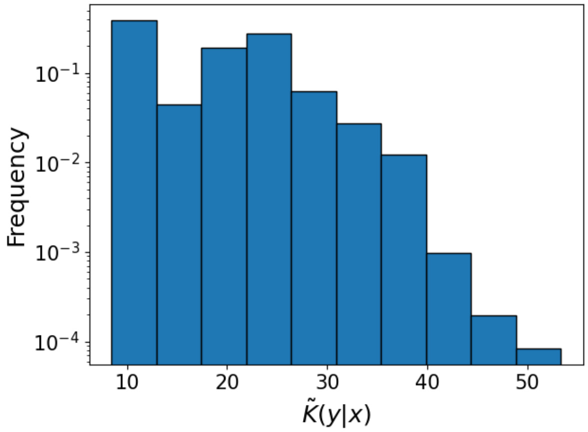


(f)

FIG. II.3. **Protein secondary structure transition probabilities for the CDR H3 hypervariable region in the antibody with PDB entry 4DQO [8] and length $L = 30$ amino acids, $x =$CCCCCCCCCCCCCEEEECCCCEEEEEECCC.** (a) Table illustrating the highest-probability SS for each conditional complexity value. The starting phenotype $x$ is marked with an asterisk (*). (b) The predicted upper bound (red) and fitted bound (black) are similar. (c) The unconditional complexity $\tilde{K}(y)$ does not predict the transition probabilities well. (d) $\tilde{K}(y|x) \approx \tilde{K}(y)$ for most $y$, leading to a positive linear correlation between values. (e) Ratios of probabilities correlate strongly with differences in conditional complexity. (f) The histogram of conditional complexity values shows a bias towards simpler proteins with lower complexity values.

enough, and e.g. ignoring $O(1)$ terms is typically not a problem. Hence, even though there is some mystery in why our predictions work, just like in some areas of computer science, mathematics, and physics, this does not negate the usefulness of these predictions.

Despite not fully understanding the success of using AIT-inspired arguments, several lines of reasoning can help us understand why they are in fact applicable, at least approximately. Firstly, although technically uncomputable, Kolmogorov complexity is fundamentally merely a measure of the size in bits of the compressed version of a data object. Hence, in many situations complexity can be approximated by standard compression algorithms (see below for more on this). Second, we stress that the upper bound derived here is specifically relevant in the practical real-world (i.e. computable, non-universal Turing machine) setting. Relatedly, Calude et al. [14] have developed an AIT for finite-state machines, suggesting that many fundamental ideas and results from AIT need not only apply to universal Turing machines. Third, the inverse connection between probability and complexity is quite intuitive, and is not therefore only expected in the limited setting in which it is studied in AIT. Fourth, many of the fundamental ideas of AIT do not depend on UTMs and uncomputable quantities, but have analogs in computable settings. For example, the fundamental fact that most sequences are incompressible results from basic counting principles, and the fact that short programs appear with higher probability in many prefix codes holds outside of UTM settings. Fifth, because a UTM can simulate any computable function, we can always bound the behaviour of a computable function with known limits of the behaviour of a UTM. In addition, Kolmogorov complexity can be bounded by lossless compression algorithms.

These points motivate using AIT as a theoretical framework to guide the derivation of mathematical predictions for real-world systems, which we might call AIT-inspired arguments.

##### B. Can we approximate Kolmogorov complexity?

Kolmogorov complexity is an uncomputable quantity, meaning that it is not possible even in principle to calculate the complexity of any arbitrary given string. However, the complexity can be bounded above. It is quite common in applications of AIT to use standard lossless compression algorithms to approximate Kolmogorov complexity [15], such as gzip of Lempel-Ziv methods for compression. Clearly these approximations have significant limitations, and cannot handle pseudorandom or chaotic patterns (see for example Cosma Shalitz (`http://bactra.org/notebooks/cep-gzip.html`) on this).

On the other hand, although technically uncomputable, Kolmogorov complexity is fundamentally merely a measure of the size in bits of the compressed version of a data object. Hence, Vitanyi [16, 17] points out that because naturally generated data is unlikely to contain pseudorandom complexities like $\pi$, the true complexity is unlikely to be much shorter than that achievable by every-day compressors. Therefore, how well standard compression algorithms work as approximations depends on the type of patterns which the system under study generates. If the system only has the computational capacity of a finite state transducer (or similar), then we can expect standard compressors to make reasonable complexity estimates. If the system has high computational capacity, or is expected to produce many pseudorandom patterns, then we can expect standard compressors not to make reasonable complexity estimates. For more on this discussion, see Ref. [16] and Ref. [18] for work on short program estimates via short lists of candidates with short programs.

##### C. Some conditions for observing simplicity bias

A full understanding of exactly which systems will, and will not, show simplicity bias (SB) is still lacking, but the phenomenon is expected to appear in a wide class of input-output maps, under fairly general conditions.

Some of these conditions were suggested in Ref. [1], including (1) that the number of inputs should be much larger than the number of outputs, (2) the number of outputs should be large, and (3) that the map should be 'simple' (technically of $O(1)$ complexity) to prevent the map itself from dominating over inputs in defining output patterns. Indeed, if an arbitrarily complex map was permitted, outputs could have arbitrary complexities and probabilities, and thereby remove any connection between probability and complexity. Finally (4), because many AIT applications rely on approximations of Kolmogorov complexity via standard lossless compression algorithms [2, 19] (but see [20, 21] for a fundamentally different approach), another proposed condition is that the map should not generate pseudorandom outputs like $\pi = 3.1415...$, which standard compressors cannot handle effectively. The presence of such outputs may yield high-probability outputs which appear 'complex' hence apparently violating SB, but which are in fact simple.

The ways in which SB differs from Levin's coding theorem include that it does not assume UTMs, uses approximations of complexities, and for many outputs $P(x) \ll 2^{-K(x)}$. Hence, the abundance of low-complexity, low-probability outputs [11, 22] is a signature of SB. Further, the upper bound was intended to apply to more general notions of complexity such as the number of stems on an RNA molecule structure, not strictly Kolmogorov complexity.

##### D. Simplicity bias: conditional form derivation

Just as for the original coding theorem, the conditional coding theorem in Eq. (6) in the main text cannot be directly applied to practical real-world systems, such as making estimates for phenotype transition probabilities. So we here derive a conditional form of the simplicity bias equation, which we subsequently apply to phenotype transition probabilities.

In our finite and computable setting we can proceed with the following argument: Given $x$ and the GP map $f$, we can enumerate genotypes and record those that map onto $x$, i.e. we can generate the NS of $x$. Next, we make a list $\mu_1(x)$ consisting of all possible 1-point mutations of the sequences in the NS of $x$. Denote the number of entries in the list $\mu_1(x)$ with $|\mu_1(x)|$. The value of $|\mu_1(x)|$ is related to the size of the genotype alphabet $\alpha$ and the length $L$ of the genotype; specifically $|\mu_1(x)|$ will be $(\alpha-1)L$ multiplied by the number of genotypes in the NS of $x$. Notice that $\mu_1(x)$ may have repeated sequences. For example, if the NS of $x$ was made up of the binary strings [001,100] then $\mu_1 =$ [101,011,000, 000,110,101] and so $|\mu_1(x)| = 6$, even though the number of unique sequences in $\mu_1(x)$ is only 4. On the other hand, for the NS given by [001,000], we have $|\mu_1(x)| = 6$ and also 6 unique members.

Next, we enumerate all genotypes in the $\mu_1(x)$ and use the map to make a list of length $|\mu_1(x)|$ of the corresponding phenotypes. Counting the frequencies of each phenotype in this list gives a discrete (and computable) probability distribution $P(x \to y)$ for all possible phenotypes $y$. (Note that $P(x \to y)$ may be 0 for some or many of the possible $y$, meaning that it might not be possible to transition directly from phenotype $x$ to phenotype $y$.) In this manner, given $x$ and the map $f$ we can use a Shannon-Fano code [23] to make a binary code using $\log_2(1/P(x \to y)) + O(1)$ bits to describe each $y$. (In fact this Shannon-Fano code will be a prefix code, meaning that no code forms the prefix of another code; see [23] for more details).

So we have

$$K(y|x,f) \leq \log_2(1/P(x \to y)) + O(1) \tag{S.3}$$

Rearranging yields the upper bound,

$$P(x \to y) \leq 2^{-K(y|x,f)+O(1)} \tag{S.4}$$

##### E. Comment on derivation

The derivation of the upper bound may appear somewhat circular, in that $K(y|x)$ is defined in terms of $P(x \to y)$, and then $K(y|x)$ is used to predict the value of $P(x \to y)$ itself. The reason that this is done is due to a fundamental notion of AIT that complexity is largely a property of an object itself, and only fairly weakly dependent on the choice of description method. This notion is formalised in the invariance theorem, which says that two UTMs will agree on the complexity of a string $y$ up to an additive constant. Hence the logic of the derivation is that *if* string $y$ can be described in a short way according to the transition probability distribution, then $y$ can also be expected to have a short description via a different complexity measure, such as a real-world compression algorithm.

Part of the 'magic' of AIT and what forms the basis of our probability predictions is utilising the approximate independence of complexity estimates. If $\tilde{K}(y|x)$ was defined in terms of $P(x \to y)$ and then used to predict $P(x \to y)$ then this would be a trivial prediction, and circular. But bounding $\tilde{K}(y|x)$ in terms of $P(x \to y)$ and then estimating $\tilde{K}(y|x)$ directly via a different method (e.g., lossless compression) yields a non-circular and valuable prediction.

##### F. Complexity of the GP map

The complexity of the GP map $f$ is an important quantity, which we discuss in this section.

If genotypes are randomly assigned to phenotypes, then the map $f$ will contain a large amount of information due to literally storing all the genotype assignments. Because there are $N_p^{N_g}$ different ways to assign $N_g$ genotypes to $N_p$ phenotypes (assuming for simplicity that phenotypes do not necessarily have to have at least one genotype assigned), this means that we can estimate the Kolmogorov complexity $K(f)$ of the map $f$ if genotypes are randomly assigned to phenotypes: $K(f) \approx \log_2(N_p^{N_g}) = N_g \log_2(N_p)$, which is typically a very large number, due to $N_g$ typically being exponential in $L$. To give a reference point, note that typical phenotypes $x$ have $K(x) \approx \log_2(N_p)$, which is far smaller. Because of the large size of $K(f)$, under the assumption of random assignment, the quantity $K(y|x,f)$ in Eq. (S.4) may be very different to $K(y|x)$ and hence the probability $P(x \to y)$ may have little relation to the complexities of $x$, $y$ or their conditional complexity. In this case, predicting $P(x \to y)$ just from these complexities will not be feasible. More generally, if the map $f$ is allowed to have arbitrary complexity values, then $f$ could be chosen such that $P(x \to y)$ takes arbitrary values, and hence will be very hard to predict.

Fortunately, many GP maps are not random, but in fact have simple fixed rule-sets for determining how genotypes are assigned to phenotypes [1]. For example, there is a fixed-size algorithm for computationally predicting an RNA SS from a given nucleotide sequence. This algorithm does not depend on the size $L$ of the sequence. The algorithm may not appear to be 'simple' in the everyday sense of word due to the various biophysical rules and constants that underlie it, but in terms of information content we can write $K(f) = O(1)$, and hence it is 'simple' in this sense.

Note that for many maps the complexity sits between the extreme values of a completely random assignment and an $O(1)$ complexity map. Presumably, the closer the map complexity is to $O(1)$, the stronger the connection between $P(x \rightarrow y)$ and $2^{-K(y|x)}$. This reasoning would predict that Eq. (10) in the main text applies approximately to any 'low'-complexity GP maps, not specifically $O(1)$ maps. We leave this to future work to explore.

##### G. Size of the genotype alphabet and number of mutations

For the bound of Eq. (10) in the main text to have stronger predictive value on point mutations, we suggest that the size of the genotype alphabet $\alpha$ should be small. This is not a very onerous condition, and is in fact quite naturally satisfied. For RNA, $\alpha = 4$ and even for protein sequences there are 20 normally occurring amino acids, so that $\alpha = 20$, while $L$ in natural RNA and proteins is much larger than these small values.

If $\alpha$ is large and $L$ is very small, then even one single point mutation will significantly randomise the genotype and possibly the correlation between the original genotype $g$ and the mutated version $g'$ may be reduced, making the relationship between phenotypes $x$ and $y$ comparatively weak. On the other hand, if $\alpha$ is small and $L$ is large (the more common case) then a single mutation will not alter the genotype strongly, so that $g$ and $g'$ will share a lot of information and be highly correlated. Hence we can expect $x$ and $y$ to be related, and follow Eq. (10) in the main text.

Additionally, for the bound to have predictive value, the number of mutations should be small: With 1 or a small number of mutations, the correlations between genotypes remains higher, but for more and more mutations, the correlation becomes very weak, and the conditional complexity becomes irrelevant.

##### H. When is $P(y)$ a good predictor of $P(x \rightarrow y)$?

From AIT we know that almost all pairs of phenotypes $x$ and $y$ share almost no information, in other words $K(x,y) \approx K(x) + K(y)$, so that

$$K(y|x) \approx K(y) \tag{S.5}$$

From this we can infer that for almost all pairs of phenotypes $x$ and $y$ the conditional complexity $K(y|x)$ in Eq. (S.4) in the SI can be replaced with just $K(y)$, and so the equation becomes

$$P(x \rightarrow y) \lesssim 2^{-K(y)} \tag{S.6}$$

for almost all $y$.

The preceding argument suggests that for most outputs $y$, the phenotype $x$ may be largely irrelevant in estimating the probabilities $P(x \rightarrow y)$. However, this statement comes with the caveat that nearly all the probability mass is likely to be associated to only a small fraction of the possible outputs, those for which $K(y|x)$ is low. In other words, with high probability $2^{-K(y)}$ may not be a good estimator of $P(x \rightarrow y)$, even if it is for most outputs $y$. Indeed, when $K(y|x) \approx 0$ but $K(y) \gg 0$, meaning that $y$ is complex and very similar to $x$, we might expect

$$P(x \rightarrow y) \gg 2^{-K(y)+O(1)} \geq P(y) \tag{S.7}$$

and hence $P(y)$ is not a good approximation of $P(x \rightarrow y)$. Potentially these types of phenotypes with $K(y|x) \approx 0$ may absorb much of the probability mass. For complex outputs $x$, even though they are rare, it could be that the majority of the $y$ for which $P(x \rightarrow y)$ is non-zero are transitions of this type.

Another important case in which conditioning on $x$ does not have much effect on the complexity or probability is the case that $x$ is very simple. If $x$ is very simple, then $K(x) \approx 0$ and $K(y|x) \approx K(y)$ for any $y$, and so we can also predict that Eq. (S.6) in the SI will hold when $x$ is very simple:

$$K(x) \approx 0 \Rightarrow K(y|x) \approx K(y) \Rightarrow P(x \rightarrow y) \lesssim 2^{-K(y)} \tag{S.8}$$

This case is important because in maps with simplicity bias, very simple outputs may be the most likely (and also common in biology [12]), hence this type of transition is likely to be quite common. However, for more complex $x$ it may be that $\tilde{K}(y|x)$ and $\tilde{K}(y)$ are different with high probability.

Whether Eq. (S.6) in the SI holds due to the fact that almost all pairs $x$ and $y$ share no information, or due to the fact that $x$ is very simple, it is interesting to further make the rough approximation that $2^{-K(y)}$ is a good estimator of $P(y)$, then this suggests that for most phenotypes $y$, we might expect

$$P(x \rightarrow y) \lesssim P(y) \tag{S.9}$$

to hold roughly, which almost recovers the null model of $P(x \rightarrow y) \approx P(y)$ described in Eq. (2) of the main text.

It is noteworthy that when Greenbury et al [24] investigated transition probabilities and concluded that $P(x \to x) \approx P(y)$ is a good estimator, or $\phi_{qp} \approx f_p$ in their terminology, they chose the phenotypes of the highest or second-highest probability as the starting phenotype $x$. For a map with simplicity bias these high-probability phenotypes would be among the simplest ones and have $K(x) \approx 0$, which is precisely the scenario in which we expect $P(x \to y)$ to be most closely related to $P(y)$. It is therefore possible that this simple prediction will break down when the starting phenotype $x$ has a lower probability, or similarly if $x$ has high complexity.

#### IV. SUPPLEMENTAL FIGURES

Here we show some extra figures, each of which shows qualitatively the same patterns as in those from the main text.

Plots for an RNA of length $L = 40$ nucleotides are shown in SI Figure II.2. Although based on a different starting phenotype, these plots look qualitatively very similar to the RNA figure in the main text.

SI Figure II.3 contains plots for the SS of a CDR H3 hypervariable region from the HIV-1-neutralizing antibody PG16 antibody with PDB entry 4DQO [8] and CDR H3 length $L = 30$. However, the SS prediction algorithm Porter 5 does not predict the original starting protein SS well (accuracy only 36% when compared with the PDB SS data [6]) and hence the probabilities and structures underlying the figure probably do not reflect the true ones that would be obtained via in vitro mutational analysis, if this were performed.

---


[1] Kamaludin Dingle, Chico Q Camargo, and Ard A Louis. Input–output maps are strongly biased towards simple outputs. *Nature communications*, 9(1):761, 2018.

[2] A. Lempel and J. Ziv. On the complexity of finite sequences. *Information Theory, IEEE Transactions on*, 22(1):75–81, 1976.

[3] Ronny Lorenz, Stephan H Bernhart, Christian Höner Zu Siederdissen, Hakim Tafer, Christoph Flamm, Peter F Stadler, and Ivo L Hofacker. Viennarna package 2.0. *Algorithms for molecular biology*, 6(1):26, 2011.

[4] Marcel Weiß and Sebastian E Ahnert. Using small samples to estimate neutral component size and robustness in the genotype–phenotype map of rna secondary structure. *Journal of the Royal Society Interface*, 17(166):20190784, 2020.

[5] Mirko Torrisi, Manaz Kaleel, and Gianluca Pollastri. Deeper profiles and cascaded recurrent and convolutional neural networks for state-of-the-art protein secondary structure prediction. *Scientific reports*, 9(1):1–12, 2019.

[6] D. Pinto, D., Park, Y.J., Beltramello, M., Walls, A.C., Tortorici, M.A., Bianchi, S., Jaconi, S., Culap, K., Zatta, F., Marco, A.D., Peter, A., Guarino, B., Spreafico, R., Cameroni, E., Case, J.B., Chen, R.E., Havenar-Daughton, C., Snell, G., Telenti, A., Vi. Structural and functional analysis of a potent sarbecovirus neutralizing antibody, DOI: 10.2210/pdb6WS6/pdb, 2020.

[7] Dora Pinto, Young Jun Park, Martina Beltramello, Alexandra C. Walls, M. Alejandra Tortorici, Siro Bianchi, Stefano Jaconi, Katja Culap, Fabrizia Zatta, Anna De Marco, Alessia Peter, Barbara Guarino, Roberto Spreafico, Elisabetta Cameroni, James Brett Case, Rita E. Chen, Colin Havenar-Daughton, Gyorgy Snell, Amalio Telenti, Herbert W. Virgin, Antonio Lanzavecchia, Michael S. Diamond, Katja Fink, David Veesler, and Davide Corti. Cross-neutralization of SARS-CoV-2 by a human monoclonal SARS-CoV antibody. *Nature*, 583(7815):290–295, 2020.

[8] P.D. Pancera, M., McLellan, J.S., Kwong. Crystal Structure of PG16 Fab in Complex with V1V2 Region from HIV-1 strain ZM109, DOI: 10.2210/pdb4DQO/pdb, 2013.

[9] Marie Pancera, Syed Shahzad-ul hussan, Nicole A Doria-rose, Jason S Mclellan, Robert T Bailer, Kaifan Dai, Sandra Loesgen, Mark K Louder, Ryan P Staupe, Yongping Yang, Baoshan Zhang, Robert Parks, Joshua Eudailey, Krissey E Lloyd, Julie Blinn, S Munir Alam, Barton F Haynes, Mohammed N Amin, Lai-xi Wang, Dennis R Burton, Wayne C Koff, Gary J Nabel, John R Mascola, Carole A Bewley, and Peter D Kwong. Structural basis for diverse N-glycan recognition by HIV-1 – neutralizing V1 – V2 – directed antibody PG16. *Nat. Publ. Gr.*, 20(7), 2013.

[10] James Dunbar, Konrad Krawczyk, Jinwoo Leem, Terry Baker, Angelika Fuchs, Guy Georges, Jiye Shi, and Charlotte M Deane. SAbDab : the structural antibody database. *Nucleic Acids Res.*, 42(November 2013):1140–1146, 2014.

[11] Kamaludin Dingle, Guillermo Valle Pérez, and Ard A Louis. Generic predictions of output probability based on complexities of inputs and outputs. *Scientific reports*, 10(1):1–9, 2020.

[12] Iain G Johnston, Kamaludin Dingle, Sam F Greenbury, Chico Q Camargo, Jonathan PK Doye, Sebastian E Ahnert, and Ard A Louis. Symmetry and simplicity spontaneously emerge from the algorithmic nature of evolution. *Proceedings of the National Academy of Sciences*, 119(11):e2113883119, 2022.

[13] Scott Aaronson. Why philosophers should care about computational complexity. *Computability: Turing, Gödel, Church, and Beyond*, 261:327, 2013.

[14] Cristian S Calude, Kai Salomaa, and Tania K Roblot. Finite state complexity. *Theoretical Computer Science*, 412(41):5668–5677, 2011.

[15] M. Li and P.M.B. Vitanyi. *An introduction to Kolmogorov complexity and its applications*. Springer-Verlag New York Inc, 2008.

[16] Paul Vitányi. How incomputable is kolmogorov complexity? *Entropy*, 22(4):408, 2020.

[17] Paul MB Vitányi. Similarity and denoising. *Philosophical Transactions of the Royal Society A: Mathematical, Physical and Engineering Sciences*, 371(1984), 2013.

[18] Jason Teutsch and Marius Zimand. A brief on short descriptions. *SIGACT News*, 47(1):42–67, March 2016.

[19] Jacob Ziv and Abraham Lempel. A universal algorithm for sequential data compression. *IEEE Transactions on information theory*, 23(3):337–343, 1977.

[20] J.P. Delahaye and H. Zenil. Numerical evaluation of algorithmic complexity for short strings: A glance into the innermost structure of algorithmic randomness. *Appl. Math. Comput.*, 219:63–77, 2012.
[21] Fernando Soler-Toscano, Hector Zenil, Jean-Paul Delahaye, and Nicolas Gauvrit. Calculating Kolmogorov complexity from the output frequency distributions of small Turing machines. *PloS one*, 9(5):e96223, 2014.
[22] Mohamed Alaskandarani and Kamaludin Dingle. Low complexity, low probability patterns and consequences for algorithmic probability applications. *arXiv preprint arXiv:2207.12251*, 2022.
[23] TM Cover and J.A. Thomas. *Elements of information theory.* John Wiley and Sons, 2006.
[24] Sam F Greenbury, Steffen Schaper, Sebastian E Ahnert, and Ard A Louis. Genetic correlations greatly increase mutational robustness and can both reduce and enhance evolvability. *PLoS computational biology*, 12(3):e1004773, 2016.
